# ADT-030, a novel PDE10 inhibitor, demonstrates potent antitumor activity in pancreatic ductal adenocarcinoma

**DOI:** 10.64898/2026.02.11.705411

**Authors:** Dhana Sekhar Reddy Bandi, Ganji Purnachandra Nagaraju, Sujith Sarvesh, Junwei Wang, Jeremy B. Foote, Adam B. Keeton, Xi Chen, Veronica Ramirez-Alcantara, Thomas Holmes, Mehmet Akce, Anju Singh, Craig M. Powell, Santosh Kumar Behera, Asfar S. Azmi, Elmar Nurmemmedov, Ivan Babic, Gregory S. Gorman, Lori Coward, Donald J. Buchsbaum, Yulia Y. Maxuitenko, Gary A. Piazza, Bassel F. El-Rayes

## Abstract

Phosphodiesterase 10 (PDE10) has been noted to be highly expressed in multiple types of cancer and is crucial for the growth and maintenance of cancer cells found in colon, lung, and ovarian cancers. Here, we studied a novel orally bioavailable PDE10 inhibitor, ADT-030, and found that it potently inhibits the proliferation and clonogenicity of KRAS-mutant pancreatic ductal adenocarcinoma (PDAC) cells at levels that block recombinant PDE10. ADT-030 also inhibited PDAC cell motility and triggered G2/M cell cycle halt and programmed cell death. These impacts were facilitated by raised cAMP/cGMP levels, activation of PKA/PKG, reduced β-catenin and RAS signaling. Notably, ADT-030 diminished the proliferation of PDAC cells with KRAS^G12D^ and KRAS^G12C^ mutations that were resistant to both allele-specific and pan-RAS inhibitors. When administered orally, ADT-030 markedly decreased tumor growth, lowered the metastasis to the lungs and liver, and enhanced survival rates without causing systemic toxicity in both syngeneic and patient-derived xenograft (PDX) models of PDAC. ADT-030 also increased chemotherapy response in orthotopic PDAC models. Immune phenotyping and single-cell RNA sequencing revealed remodeling of the tumor microenvironment by ADT-030 with a more favorable anti-tumor immune profile. The findings suggest that ADT-030 holds promise as a potential drug development candidate for treating KRAS-mutant PDAC by simultaneously targeting key oncogenic signaling pathways, resulting in tumor-intrinsic and immunomodulatory effects.

## Introduction

Pancreatic ductal adenocarcinoma (PDAC) is the fourth most frequent reason of cancer-related mortality in the U.S., with a five-year survival of under 13% ^1^. Survival of patients with metastatic PDAC remains poor even with the current chemotherapeutic treatments such as FOLFIRINOX or gemcitabine and nab-paclitaxel ^2^. The aggressive nature of PDAC is mainly attributed to activation of multiple compensatory signaling pathways driving proliferation and survival, along with a hypoxic microenvironment driven by dense desmoplastic stroma and decreased vascular perfusion ^3–7^. Although both allele-specific KRAS and pan-RAS inhibitors have demonstrated promising activity in patients with PDAC during clinical trials, the expansion of resistance remnants a main challenge and highlights the need to identify new therapeutic targets and agents with more durable anticancer activity and capacity to overcome resistance ^8–10^. A better comprehension of the complexity of oncogenic signaling, the consequence of stroma, and the function of immune evasion in PDAC advancement is critical for the development of more effective target-directed drugs for the treatment of PDAC ^11^.

Mutations in the KRAS gene have been noted in about 90% of PDAC patients, with the common appearing at codon 12 (KRAS^G12D^, KRAS^G12V^, and KRAS^G12C^) ^12^. These mutations lead to constant activation of downstream pathways, involving RAS/RAF/MEK and PI3K/AKT/mTOR signaling, thereby encouraging the proliferation, survival, and metabolic reprogramming of pancreatic ductal adenocarcinoma (PDAC). ^13^. Although the mutation frequency of *CTNNB1* is relatively low in PDAC, β-catenin signaling is aberrantly activated from WNT overexpression, which, along with KRAS mutations, contributes to the aggressive performance of PDAC ^6^. KRAS has been reported to form a complex with β-catenin, modulating the phosphorylation of the transcription factor TCF4 and leading to crosstalk between these oncogenic signaling pathways ^14,15^. In addition, both β-catenin and RAS signaling have been reported to be activated with gemcitabine treatment, suggesting that these pathways cooperate to cause therapy resistance in PDAC ^16^.

Recent studies have also conveyed in a metastatic mouse model of colorectal cancer that Wnt activation of Lgr5+ stem cell-like can drive resistance to RAS inhibitors ^17^. Given the interactions between RAS and Wnt/β-catenin signaling in PDAC, a strategy that targets both pathways with a single inhibitor could offer a more robust and durable therapeutic approach. Emerging evidence also suggests that simultaneous inhibition of multiple oncogenic pathways not only increases the potential for efficacy of RAS-specific inhibitors but also reduces the potential for resistance by overcoming compensatory signaling mechanisms ^18,19^. KRAS-mutated tumors can utilize β-catenin signaling to maintain a stem-like, immune-depleted niche within the tumor immune microenvironment (TiME), resulting in relapses following chemotherapy. Targeting both pathways with a single inhibitor could also sensitize cancer cells to undergo apoptosis while modulating the TiME to favor immune activation, to escape resistance ^20^.

Phosphodiesterase (PDE) isoenzymes hydrolyze and inactivate the second messengers, cyclic adenosine monophosphate (cAMP) and cyclic guanosine monophosphate (cGMP) ^21^. Although understudied, PDEs have been reported to play a role in the initiation and progression of PDAC and other cancers ^22,23^. Notably, the cAMP/PKA and cGMP/PKG signaling axes have been reported to suppress MAPK signaling downstream from KRAS ^24,25^. Several PDE isozyme families, most notably PDE4, PDE5, and PDE10, have been investigated as anti-cancer targets affecting cancer cell proliferation, survival, and immune responses ^23,26^. Recently, isozyme-specific inhibitors and gene silencing of the dual cAMP/cGMP-degrading PDE10 isozyme have been stated to selectively inhibit the proliferation and induce apoptosis of cells from colon, lung, and ovarian cancers by disrupting RAS and WNT/β-catenin signaling ^27–30^. These findings established the basis for our studies of ADT-030, a novel PDE10 inhibitor. Unlike PDE10 inhibitors developed for CNS disorders, oral administration of ADT-030 achieved high and sustained systemic levels in mice sufficient to block tumor growth. We report here that the mechanism involves PDE10 inhibition and activation of protein kinases to block both RAS and β-catenin signaling, ensuing in the destruction of PDAC tumor growth and metastasis, as well as activation of antitumor immunity. By targeting PDE10, ADT-030 shows the potential to evade compensatory mechanisms of resistance that limit the efficacy of other RAS inhibitors and with the potential to mitigate toxicities associated with Wnt/β-catenin inhibitors. The findings encourage IND-enabling studies to improve ADT-030 to clinical trials as a monotherapy or in combination with standard-of-care chemotherapy.

## Results

### PDE10 overexpression in PDAC correlates with KRAS signaling and disease progression

To initially investigate the role of PDE10 in PDAC development, we used 10x Genomics sequencing to measure PDE10 expression in normal and cancerous tissues from 229 patients across 12 different study groups. Employing sc-RNA seq data, we found that PDE10 mRNA is extensively enriched in primary tumors and metastatic lesions associated to tissues from normal donors and adjacent uninvolved tissues (**Fig. 1a**). Transcriptomic analysis of the TCGA PDAC dataset exhibited that PDE10 expression is extensively eminent in pancreatic tumors compared with normal pancreatic tissue (**Fig. 1b**). In addition, Log_2_-transformed expression values revealed a statistically significant positive correlation between PDE10 and KRAS mRNA levels (Pearson r = 0.37, p < 0.0001, **Fig. 1c**). These findings identify PDE10 as a PDAC-associated gene that is enriched within the KRAS-driven PDAC transcriptional landscape. We then tested the PDAC cell lines used in the experiments described below and found that all expressed PDE10 at comparable levels (**Fig. 1d**).

**Fig. 1:**
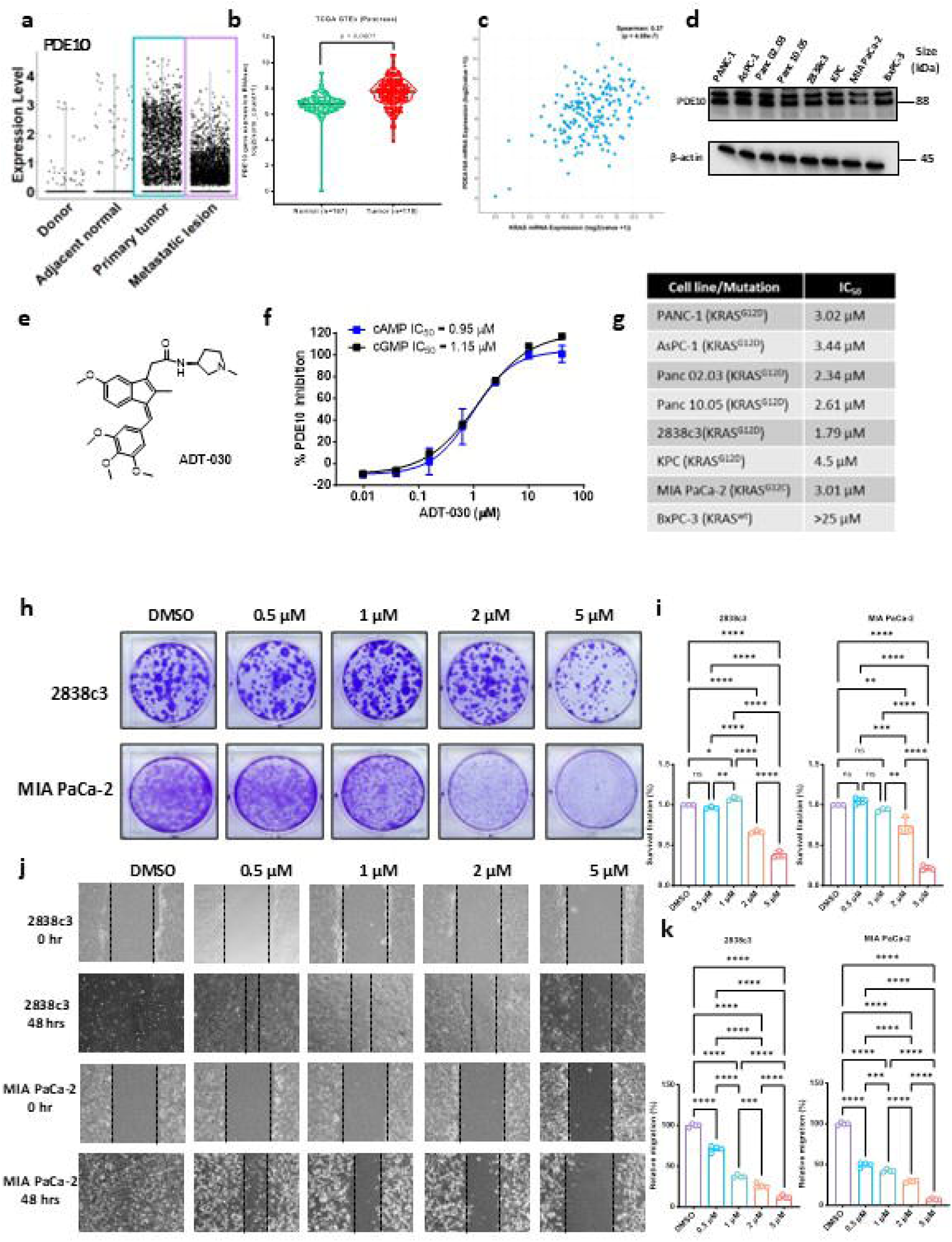
ADT-030 inhibits PDE10 at concentrations that inhibit proliferation, colony formation, and motility, while inducing apoptosis and cell cycle arrest of KRAS mutant PDAC cells. **a.** Violin plot of PDE10 expression in human donor, adjacent normal tissue, primary tumor and metastatic lesion as determined by sc-RNAseq analysis of human PDAC. **b.** The TCGA-GTEx comparison demonstrates that PDE10 mRNA expression is significantly higher in pancreatic tumors than in normal pancreatic tissue (P < 0.0001). **c.** Correlation analysis reveals a significant positive relationship between PDE10 and KRAS mRNA expression (Spearman correlation = 0.37; P = 4.08 × 10^-7^), suggesting that tumors with higher KRAS expression tend to exhibit higher PDE10 expression. **d.** Baseline expression of PDE10 across indicated PDAC cell lines. β-actin was used as a loading control. **e.** Chemical structure of ADT-030. **F.** ADT-030 inhibits the enzymatic activity of recombinant PDE10 using cAMP and cGMP as substrates. Data are expressed as mean ± SD, n = 2 samples/concentration. **g.** PDAC cell lines were treated with various concentrations of ADT-030 for 3 days, followed by determining viable cell number using MTT assays. Relative percentage cell viability was plotted with respect to vehicle (DMSO) treated cells. The table lists the IC_50_ values for the PDAC cell lines treated with ADT-030. **h-i.** The indicated PDAC cell lines were treated with various concentrations of ADT-030 for 2–4 weeks, and long-term cell survival was measured using clonogenic assays. Representative images are shown in **h** and quantification plotted in **i**. **j.** Indicated PDAC cell lines were treated with vehicle or the indicated concentrations of ADT-030, and cell migration was analyzed using cell motility assays. Representative images under indicated treatment conditions for indicated PDAC cell lines are shown. **k.** Bar diagrams are presented to show relative migration (%) from the experiment presented in **j**. Data represent the mean ± SEM of three biological replicates. ns = not significant, *p < 0.05, **p < 0.01, ***p < 0.001, ****p < 0.0001. (one-way ANOVA).

### ADT-030 inhibition and binding of PDE10

ADT-030 is an indene chemically related to the nonsteroidal anti-inflammatory drug, sulindac, designed to block cyclooxygenase (COX) inhibitory activity while targeting PDE10 (**Fig. 1e**) as previously reported for an analog, ADT-061 ^30^. The potency of ADT-030 to inhibit the enzymatic activity of recombinant PDE10 was determined by measuring cGMP and cAMP hydrolysis using a fluorescence polarization assay. ADT-030 inhibited cAMP and cGMP hydrolysis with IC_50_ values of 0.95 and 1.15 µM, correspondingly (**Fig. 1f**). Molecular modeling studies using the PDE10 (2OUN) structure were performed by induced-fit molecular dynamics and simulation interaction analysis to identify a potential binding site on PDE10 for ADT-030. An optimal GLIDE docking score of -10.3 kcal/mol was calculated with ADT-030 bound in the PDE10 catalytic domain with the tri-methoxy benzylic moiety oriented toward the deep hydrophobic region of the pocket, while the more polar substituents were oriented towards the pocket entrance (**Supplementary fig. 1a**). The amide carbonyl and neighboring heteroatoms are predicted to form hydrogen bonds with His525 through bridging water molecules (**Supplementary fig. 1b**). Hydrophobic and aromatic contacts with residues Tyr524, Leu635, Phe639, Ile692, Tyr693, Phe696, Met714, Phe729, Val733, and Ala734 help to anchor the aromatic system within the binding site. These results suggest a favorable binding for ADT-030 in the PDE10 catalytic domain, which is supported by the docking score and a network of direct and water-mediated interactions, and consistent with a competitive mechanism of enzyme inhibition as previously reported for ADT-061 ^30^.

Experiments were also performed to confirm that ADT-030 binds PDE10 in intact cells using a cellular engagement thermal stability assay. In brief, HEK-293 cells expressing PDE10 Micro-Tag were treated with ADT-030. PDE10 thermal stability was measured by Micro-Tag enzyme complementation as explained in the Materials and Methods section (**Supplementary fig. 1c-f**). The results revealed an EC_50_ of 0.9 µM for ADT-030 to bind PDE10, which paralleled its potency in inhibiting PDE10 enzymatic activity.

### ADT-030 inhibition in wild-type vs. mutant KRAS PDAC cell lines

The antiproliferative activity of ADT-030 was tested against a series of PDAC cell lines harboring various KRAS mutations, as well as wild-type RAS, by performing cell viability measurements using the MTT assay and determining potency (IC_50_) values. As shown in **Fig. 1g**, ADT-030 inhibited the proliferation of all KRAS^G12D^ and KRAS^G12C^ mutant PDAC cells tested with IC_50_ values in the low micromolar range (1.8-4.5 µM). Notably, the KRAS wild-type PDAC cell line, BxPC-3, was found to be essentially insensitive to ADT-030, suggesting that ADT-030 selectively reduces the growth of KRAS mutant PDAC cells (**Fig. 1g**). PDE10 expression was knocked down using shRNAs in AsPC-1 and PANC-1 cells, both of which harbor KRAS^G12D^ mutation (**Supplementary fig. 2a, b**). PDE10 knockdown sensitized PDAC cells to ADT-030 compared with parental cells (**Supplementary fig. 2c, d**). These findings demonstrate that PDE10 expression directly influences ADT-030 sensitivity and support PDE10 as the functional molecular target of ADT-030 in PDAC.

We next investigated whether ADT-030-induced metabolic rewiring differed between KRAS wild-type and mutant PDAC cells. Lysates from ADT-030- and DMSO-treated PANC-1 KRAS^G12D^ and KRAS wild-type BxPC-3 PDAC cells (**Supplementary fig. 3a, b**) were analyzed using the Human Metabolic Pathways Array C2. ADT-030 elicited distinct metabolic responses depending on KRAS mutational status. In KRAS-mutant cells, ADT-030 treatment resulted in downregulation of multiple metabolic enzymes involved in biosynthetic flux, redox balance, lipid metabolism, and cellular homeostasis, including ARSB, ASAH2, CA14, HAGH, ENPP2, NQO1, and PCK1 (**Supplementary fig. 3c-l**). In contrast, these same proteins were upregulated in KRAS wild-type cells following ADT-030 treatment, indicating selective suppression of a metabolic program that KRAS-mutant cells rely on for survival and proliferation. The directed downregulation of PCK1 and NQO1 enzymes is critical for gluconeogenesis-derived biosynthetic flux and redox homeostasis, suggesting that ADT-030 induces metabolic exhaustion and lethal oxidative stress, particularly in KRAS mutant cells. In addition, the suppression of pH-regulatory enzymes (CA13/14) and pro-survival lipid modulators (ASAH2/ENPP2) further supports a model of metabolic collapse (**Supplementary fig. 3c-l**). A limited subset of proteins, including EXTL3 and IDO1, exhibited divergent expression patterns, showing increased levels in mutant cells relative to wild-type controls following treatment. Together, these results present direct experimental proof that the metabolic effects of ADT-030 are influenced by KRAS mutational status, with KRAS mutant cells exhibiting heightened vulnerability to ADT-030-mediated metabolic disruption.

### ADT-030 inhibits the clonogenicity and migration of KRAS mutant PDAC cells

A mouse PDAC cell line, 2838c3 (KRAS^G12D^ mutant), and a human PDAC cell line, MIA PaCa-2 (KRAS^G12C^ mutant), were selected to further study the anti-proliferative activity of ADT-030. The long-term inhibitory effect of ADT-030 on cancer cell survival was evaluated in both PDAC cell lines by colony formation assays. ADT-030 treatment significantly reduced the number and size of colonies in both PDAC cell lines across a concentration range comparable to its potency (IC50) for inhibiting proliferation (**Fig. 1h, i**). In addition, 2838c3 and MIA PaCa-2 cells showed significant impairment in motility after treatment with ADT-030 at non-cytotoxic concentrations (**Fig. 1j, k**). Together, these results indicate that ADT-030 impedes the proliferation, colony formation, and motility of PDAC cells with KRAS^G12D^ and KRAS^G12C^ mutations within the same concentration range essential to inhibit recombinant PDE10 and to bind PDE10 in cells.

### ADT-030 triggers cell apoptosis and causes G2/M phase cell cycle arrest

To uncover the effect of ADT-030 on apoptosis and cell cycle progression, the 2838c3 and MIA PaCa-2 PDAC cells were dosed with ADT-030 for 24 and 72 hrs, respectively. Flow cytometry analysis of apoptosis as measured by Annexin V/PI staining showed that ADT-030 increased both early and late apoptotic cells at concentrations of 2 and 5 µM. In vehicle-treated 2838c3 cells, apoptotic cells comprised 1.6% of the population, whereas treatment with ADT-030 increased the percentage to 3.5% (2 µM) and ∼8% (5 µM) (**Supplementary fig. 4a, c**). For MIA PaCa-2 cells, vehicle-treated cells had 2.3% of apoptotic cells within the population, whereas ADT-030 treatment increased the number to 14.5% (2 µM) and 32% (5 µM) (**Supplementary fig. 4b, d**). Assessment of cell cycle dissemination revealed that ADT-030 treatment improved the ratio of cells arrested in the G2/M phase in both PDAC cell lines (**Supplementary fig. 4e-h**).

### ADT-030 inhibits PDE10 and activates PKA/PKG signaling

Since PDE10 inhibition by ADT-030 is expected to improve intracellular cAMP and cGMP, we measured these levels in the 2838c3 and MIA PaCa-2 PDAC cell lines following ADT-030 treatment by ELISA. Consistent with a PDE10 inhibitor, ADT-030 significantly boosted cAMP and cGMP levels in a concentration-dependent manner. Notably, the effect was apparent at concentrations that paralleled the range effective for inhibiting recombinant PDE10, inhibiting proliferation of both PDAC cells, and causing cell cycle arrest and apoptosis (**Fig. 2a-d**). To witness whether increased cyclic nucleotide levels by ADT-030 activated downstream protein kinases, PKA and PKG, in 2838c3 and MIA PaCa-2 cells, we measured the phosphorylation of VASP (vasodilator-stimulated phosphoprotein), a known substrate for PKA and PKG ^31^. Consistent with a PDE10 inhibitor, ADT-030 increased VASP phosphorylation in both PDAC cell lines (**Fig. 2e, f)**, demonstrating that ADT-030 elevated intracellular cAMP and cGMP levels, thereby activating PKA and PKG. Finally, it should be noted that ADT-030 treatment did not alter the expression of PDE10 (**Fig. 2e**) or other cGMP and cAMP-degrading PDE isozymes, PDE3 or PDE4, respectively (**Supplementary fig. 5a**). We also determined if ADT-030 can activate canonical downstream signaling from PKA/PKG activation commonly reported in normal cells. Treatment of 2836c3 and MIA PaCa-2 cells with ADT-030 did not impact the level of activated phospho-CREB, VEGF-A, and Bcl-2 (**Supplementary fig. 5b, c**), suggesting that the activation of PKA and/or PKG by ADT-030 in PDAC cells may involve downstream targets unique to cancer cells.

**Fig. 2.**
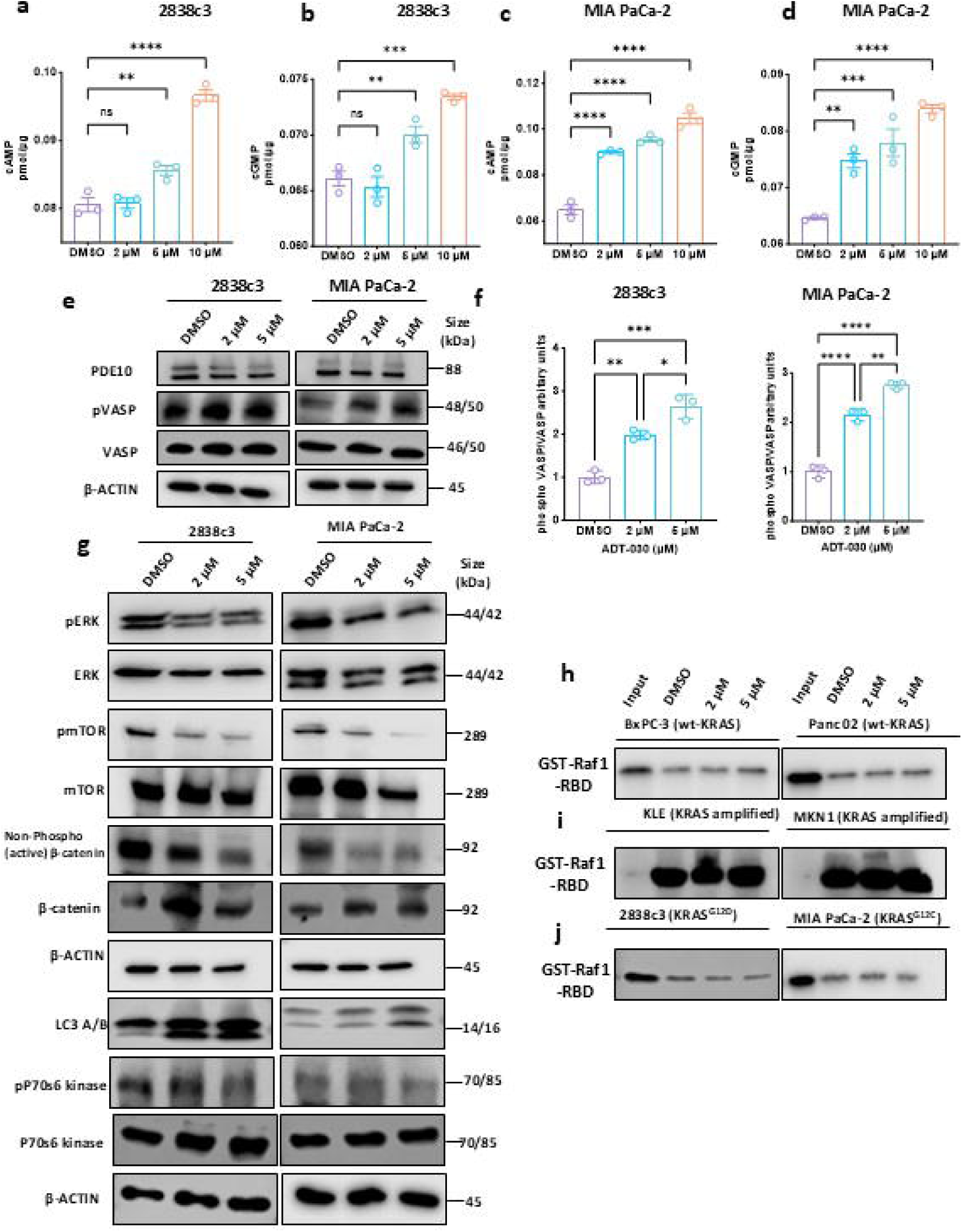
ADT-030 blocks PDE10 and activates PKA/PKG to reduce β-catenin levels and inhibit RAS signaling. **a-d.** ADT-030 increases cyclic nucleotide levels in a concentration-dependent manner in 2838c3 (**a-b**) and MIA PaCa-2 cells (**c-d**). **e-f.** Treatment with ADT-030 for 4 hrs did not affect the expression of PDE10 but induced VASP phosphorylation at serine 157 (PKA site) and serine 239 (PKG site) in a concentration-dependent manner in 2838c3 and MIA PaCa-2 cells. β-actin was used as a loading control. **g.** ADT-030 decreased phosphorylation of ERK, mTOR, and levels of active (oncogenic) β-catenin in 2838c3 and MIA PaCa-2 cells in a concentration-dependent manner. β-actin was used as a loading control. **h-j.** The indicated cell lines were treated with increasing concentrations of ADT-030 for 24 hrs, and RAS pulldown was performed. Data represent the mean ± SEM of three biological replicates. ns = not significant, *p < 0.05, **p < 0.01, ***p < 0.001, and ****p < 0.0001 (one-way ANOVA).

### ADT-030 attenuates RAS and β-catenin signaling to promote apoptosis

Previous reports suggest that known PDE10 inhibitors and PKG activation can phosphorylate the oncogenic pool of β-catenin in cell lines from various cancers ^27–30^. We therefore performed Western blot analysis to measure levels of the unphosphorylated (stable) form of β-catenin, which signifies the oncogenic pool required for TCF/LEF transcriptional activity. ADT-030 treatment significantly decreased levels of the unphosphorylated form of β-catenin in 2838c3 and MIA PaCa-2 PDAC cell lines (**Fig. 2g**). Previous research also suggested that PDE10 inhibitors and activation of PKG in lung and ovarian cancers suppress MAPK and AKT signaling ^27,29^. Hence, we determined whether ADT-030 has a similar effect in PDAC cells by measuring phosphorylated levels of ERK (pERK) and mTOR (pmTOR) within the MAPK and AKT signaling nodes, respectively. ADT-030 reduced pERK levels at its triggering phosphorylation sites, Thr^202^ and Tyr^204^, as well as pmTOR at its activation site (Ser^2448^) (**Fig. 2g**). These outcomes propose that the PDE10 repressing activity of ADT-030 can simultaneously suppress RAS and β-catenin signaling in PDAC cells.

To further study the influence of ADT-030 on RAS signaling, RAS-GTP pulldown assays were performed to measure activated RAS levels following the treatment of PDAC cells with ADT-030 for 24 hrs at concentrations of 2 and 5 µM. ADT-030 treatment did not reduce activated RAS levels in KRAS wild-type BxPC-3 and Panc 02 cells, as well as in KRAS amplified KLE (endometrial adenocarcinoma) and MKN1 (gastric adenocarcinoma) cancer cell lines (**Fig. 2h, i** and **Supplementary fig. 6a-d**). Conversely, ADT-030 reduced activated RAS levels in 2838c3 cells expressing KRAS^G12D^ and MIA PaCa-2 cells expressing KRAS^G12C^ mutations at concentrations effective for inhibiting proliferation and PDE10 (**Fig. 2j** and **Supplementary fig. 6e, f**). These results suggest that ADT-030 can inhibit activated RAS levels in KRAS-mutant PDAC cells but not in KRAS wild-type or KRAS-amplified cells. This may explain the selective growth-inhibitory activity of ADT-030 observed in RAS-mutated vs. RAS-wild-type PDAC cells.

We also explored the impact of ADT-030 treatment on autophagy in 2838c3 and MIA PaCa-2 PDAC cells, given previous studies reporting that RAS signaling and autophagy are interconnected and that RAS can modulate autophagy to promote tumorigenicity ^32^. Autophagy was assessed by Western blotting using LC3A/B as a marker. ADT-030 treatment increased LC3A/B expression, indicating its capacity to disrupt autophagic flux (**Fig. 2g**). The cells were also treated with ADT-030 alone or in combination with hydroxychloroquine (HCQ), a known autophagy inhibitor. ADT-030 in combination with HCQ did not increase the levels of LC3A/B, suggesting that ADT-030, like HCQ, inhibits autophagic flux (**Supplementary fig. 7a**), and is consistent with a previously reported analog of ADT-030 ^33^. Furthermore, ADT-030 treatment reduced p70S6 kinase expression in 2838C3 and MIA PaCa-2 cells, which is associated with reduced migratory capacity (**Fig. 2g**) ^34^.

### Hematologic, clinical chemistry, histopathologic, and behavioral assessment of ADT-030-treated mice

Our next objective was to evaluate the tolerance of mice to orally (gavage) administered ADT-030. Ten mice were randomly assigned to vehicle (n=5) or ADT-030 (150 mg/kg, n=5) treatment by oral gavage for two weeks. Complete blood counts (WBC, RBC, HGB, HCT, MPV, MCV, MCHC, RDW, PLT, MCH, neutrophils, lymphocytes, monocytes, eosinophils, and basophils) and serum biochemistry (albumin, ALT, ALP, amylase, total bilirubin, BUN, phosphorus, creatinine, glucose, electrolytes (calcium, sodium, and potassium), total protein, and globulin were measured following two weeks of treatment (**Supplementary fig. 8a, b**). We observed no noteworthy differences between vehicle- and ADT-030-treated mice, except for a slight reduction in total bilirubin levels in the ADT-030-treated group. In addition, gross examination of multiple organs, including lungs, liver, kidney, pancreas, heart, duodenum, colon, spleen, thymus, brain, and testis, showed no histopathological abnormalities in ADT-030-treated mice compared with vehicle-treated mice **(Fig. 3a, b)**.

**Fig. 3.**
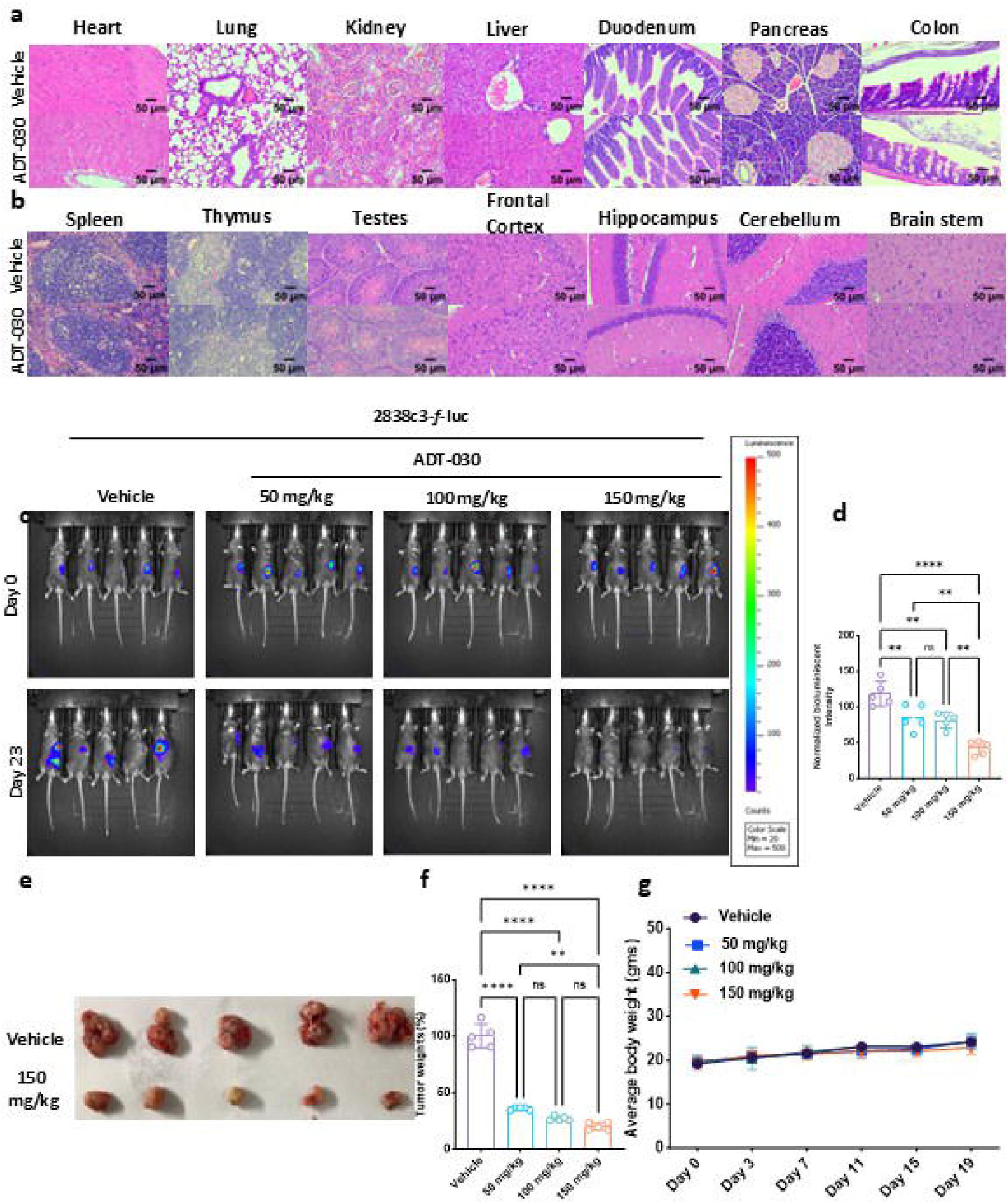
Effect of ADT-030 treatment on tumor growth and modulation of TME in 2838c3 cell-implanted C57BL/6J mice. **a-b.** Histopathological examination of vital organs (heart, lung, kidney, liver, duodenum, pancreas, colon, spleen, thymus, tests, and brain) from mice treated with ADT-030 at 150 mg/kg as assessed by HCE staining. **c.** 2838c3*-f-*luc cells were injected orthotopically into the pancreas of C57BL/6J mice. Representative bioluminescence images at the indicated time points are shown. **d.** Relative normalized whole-body bioluminescence intensities in mice under the indicated conditions (n = 5). Statistical significance was determined using one-way ANOVA. **e.** Images of the pancreatic tumors in the vehicle- and the 150 mg/kg ADT-030-treated groups at termination. **f.** Tumor weights at the end of the experiment for the indicated doses. **g.** Average body weights of mice treated with vehicle and ADT-030 (50, 100, and 150 mg/kg). Statistical significance was determined using one-way ANOVA. ns = not significant, **p < 0.01, and ****p < 0.0001.

Sedation is a well-known side effect of conventional PDE10 inhibitors that cross the blood-brain barrier (BBB) and were established for the treatment of schizophrenia and Huntington’s disease. We therefore performed an open-field locomotor test to determine if ADT-030 causes sedation. These behavioral experiments discovered no substantial changes between vehicle and ADT-030-treated mice, demonstrating that ADT-030 does not cause sedation, a common side effect from previously developed PDE10 inhibitors for CNS disorders (**Supplementary fig. 9a**).

### Pharmacokinetics and tissue distribution of ADT-030

A PK study was conducted in female C57BL/6J mice following oral gavage dosing of ADT-030 at 100 mg/kg once daily for 14 days. ADT-030 generated plasma levels that exceeded those necessary to inhibit PDE10 and PDAC cell growth *in vitro* (**Supplementary fig. 9b**). Plasma levels of ADT-030 reached a Cmax of 7 µM by 1 hr post-treatment and remained unchanged for an additional hour before decreasing by 4 hrs post-treatment to the level of 5 µM. High levels of ADT-030 were also detected in various organs (lungs, kidneys, spleen, heart, liver, ovaries, and colon) 8 hrs after administration, but low levels were measured in the brain (**Supplementary fig. 9c**). The low concentration of ADT-030 measured in the brain following oral administration likely accounts for the absence of sedation, a known side effect of conventional PDE10 inhibitors established for the treatment of CNS disorders that were designed to cross the BBB to attain high concentrations in the brain ^35^.

### ADT-030 suppresses tumor growth in the orthotopic PDAC model and reprograms the TiME

A mouse model of PDAC, involving orthotopic implantation of 2338c3-f-luc PDAC cells into the pancreas, was used to evaluate the *in vivo* antitumor activity of ADT-030. Mice that established palpable tumors one week following implantation were randomized into four groups and treated by oral gavage administration with vehicle or ADT-030 at dosages of 50, 100, and 150 mg/kg once daily 5x/week for 23 days. Tumor progression was monitored using bioluminescence imaging (**Fig. 3c**). All dosages of ADT-030 were effective, with the highest dose tested showing tumor regression in all mice in the group. Quantification of bioluminescence confirmed a substantial reduction in tumor mass in ADT-030-treated mice compared to vehicle treatment (**Fig. 3d**). Both tumor images and tumor weight measurements confirmed tumor shrinkage in ADT-030-treated groups in a dose-dependent manner compared to vehicle treatment (**Fig. 3e, f** and **Supplementary fig. 10a, b**). ADT-030 treatment did not cause apparent systemic toxicity as evidenced by no effect on body weight gain during treatment (**Fig. 3g**).

To study the immunomodulatory effects of ADT-030 relevant to PDAC, we performed multiparametric flow cytometry on orthotopic 2838c3 tumors from vehicle and ADT-030-treated mice. The analysis revealed that ADT-030 induced profound shifts in immune cell composition in the TiME, favoring a more immunostimulatory phenotype. ADT-030 treatment enhanced immune cell infiltration within the TiME, lead to in a major increase in the overall population of CD45^+^ leukocytes compared with vehicle treatment (**Supplementary fig. 11a**). Further characterization of the T-cell compartment revealed an increase in CD3^+^ T cells (**Supplementary fig. 11b**), observed with both CD4^+^ (**Supplementary fig. 11c**) and CD8^+^ T cells (**Supplementary fig. 11d**). Treatment with ADT-030 induced higher levels of several immune checkpoint markers, including CTLA-4 (**Supplementary fig. 11e**), PD-1 (**Supplementary fig. 13f**), LAG-3 (**Supplementary fig. 11g**), and TIGIT (**Supplementary fig. 11h**) in the total T cell populations. In another experiment using the KPC-f-luc model, immune checkpoint markers were differentially regulated, with a decline in CD8+ T cells and an increase in CD4^+^ T cells. These results indicate that there was a concurrent adaptive immune regulatory response, likely representative of an acute but low T cell phenotype, probably exhausted, within the TiME. In addition to modulating the T cell compartment, ADT-030 also increased NK cell infiltration within the TiME. Flow cytometry findings revealed a pronounced boost in the NK1.1^+^ cell population in the tumors of the ADT-030-treated mice compared to vehicle-treated mice, indicating enhanced activation of the innate immune system (**Supplementary fig. 11i**). In addition to augmenting NK cell numbers, increased expression of immune checkpoint receptors was measured on NK cells by ADT-030 treatment. These observations were paralleled within the CD3^+^ T cell population, where ADT-030 enhanced infiltration of CD4^+^ and CD8^+^ T cells while upregulating CTLA-4, PD-1, LAG-3, and TIGIT (**Supplementary fig. 11e-h**). These data demonstrate that ADT-030 has the capacity to broadly remodel the immune landscape, attracting both adaptive and innate effector cells into the tumor while simultaneously engaging checkpoint regulatory pathways.

### ADT-030 enhances antigen-specific T cell responses in PDAC

To further evaluate whether the anti-tumor effects of ADT-030 were accompanied by adaptive immune induction, we performed an IFN-γ ELISpot assay using splenocytes and tumor-infiltrating lymphocytes (TILs) harvested from pancreatic tumors formed by orthotopic implantation of 2838c3 cells into the mouse pancreas. We observed a noteworthy increase in the IFN-γ spot-forming cells (SFCs) in the lymphocytes from mice treated with ADT-030 compared to vehicle-treated controls (**Supplementary fig. 11j**). Upon stimulation with tumor-specific antigens (PMA+Ionomycin), lymphocytes from ADT-030-treated mice exhibited a substantial improve in IFN-γ compared to vehicle-treated mice (**Supplementary fig. 11k**). The combination of the flow cytometry and ELISpot results confirm that PDE10 inhibition by ADT-030 increased the number and the frequency of functional, cytokine-secreting effector T cells.

### ADT-030 alters RAS and β-catenin signaling in tumors

To determine the commotion of ADT-030 treatment to inhibit oncogenic signaling *in vivo*, 2838c3 tumors were harvested from mice treated with vehicle or ADT-030 and analyzed for key signaling and apoptosis markers. Western blot analysis revealed a marked decrease in pERK (**Fig. 4a, b**) and pAKT (**Fig. 4a, c**) levels with no effect on total ERK or AKT levels in tumors from ADT-030-treated mice, indicating the concurrent inhibition of the MAPK and PI3K/AKT pathways, respectively, reflective of upstream RAS inhibition. In addition, ADT-030 treatment decreased levels of the non-phosphorylated form of β-catenin, indicative of the stable pool of β-catenin driving transcription of proteins involved in oncogenesis, for example, from aberrant activation of WNT signaling (**Fig. 4a, d**). Along with these alterations in key signaling node proteins, ADT-030 treatment also increased the expression of LC3A/B, consistent with *in vitro* experiments, indicating that ADT-030 inhibits autophagic flux (**Fig. 4a, e**). An increase in cleaved PARP (Fig. 4A and 4F) and cleaved caspase 3 (Fig. 4a, g) was also observed in the ADT-030-treated group, again consistent with *in vitro* experiments showing apoptosis induction by ADT-030 treatment. Also consistent with *in vitro* experiments, ADT-030 treatment reduced activated (GTP-bound) RAS levels as measured by RAS-RBD pulldown assays (**Fig. 4h** and **Supplementary fig. 13l**). IHC showed significantly reduced expression of the proliferation marker, Ki-67, in tumors from ADT-030-treated mice, corroborating the antiproliferative activity of ADT-030 as observed *in vitro* (**Fig. 4i, m**). IF microscopy evaluation was used to analyze the treatment impact on autophagy and mesenchymal-to-epithelial transition (MET). Mice treated with ADT-030 at 150 mg/kg showed an increased level of LC3A/B in tumors, indicative of a disruption of autophagic flux (**Fig. 4j, n**). ADT-030 also reduced vimentin expression (**Fig. 4k, o**) and increased E-cadherin expression (**Fig. 4l, p**), both of which are associated with MET and indicative of transforming cancer cells into a more normal epithelial phenotype with lower invasive capability.

**Fig. 4.**
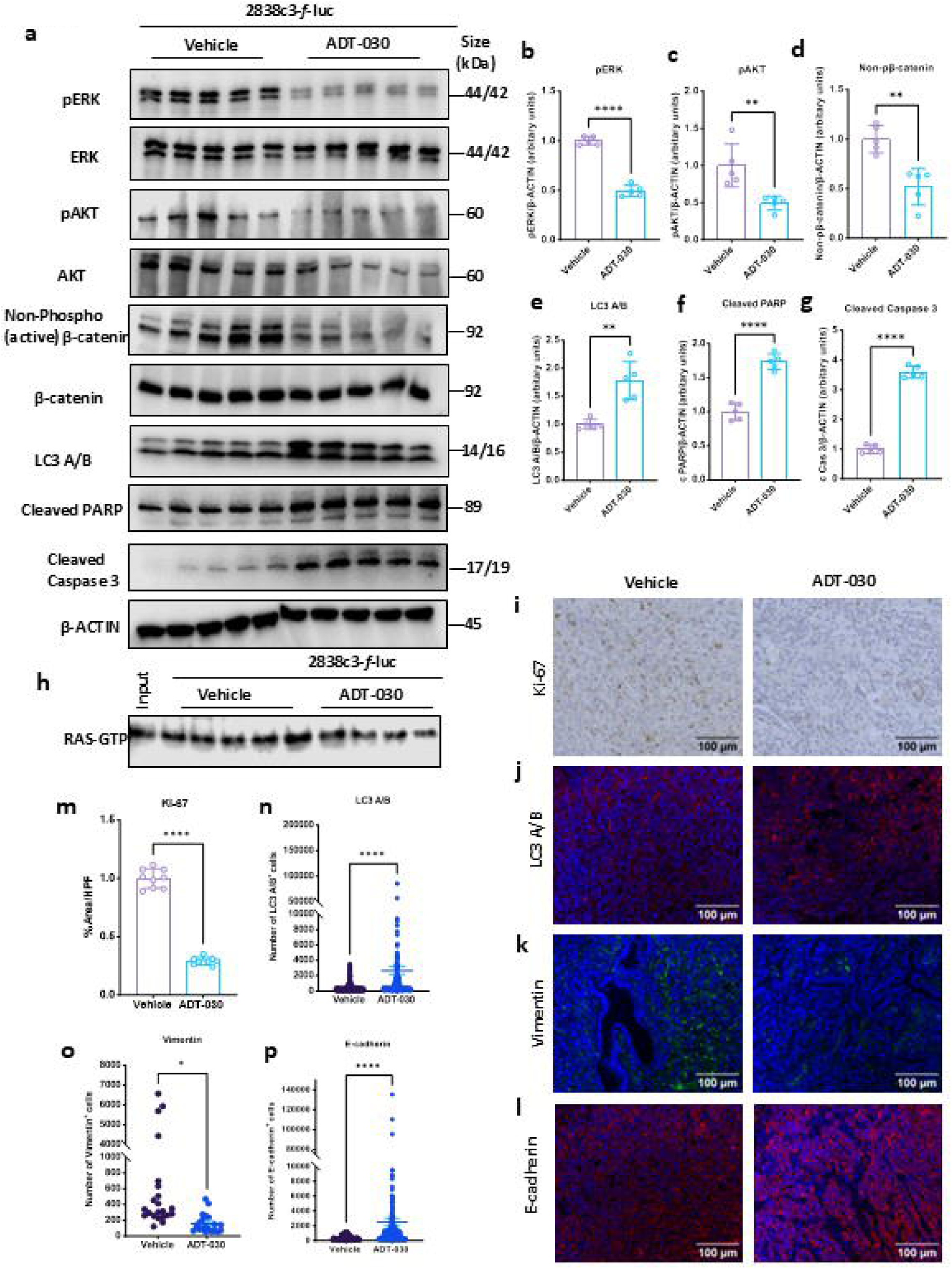
ADT-030 reduces β-catenin levels, inhibits RAS/AKT signaling, and induces autophagic cell death. **a.** Western blots showing levels of pERK, total ERK, pAKT, total AKT, non-phospho-β-catenin, total β-catenin, LC3A/B, cleaved PARP, and cleaved caspase 3 in 2838c3*-f-*luc tumors after vehicle or ADT-030 treatment. **b-g.** Bar graphs representing the quantifications of western blots from panel **a:** pERK (**b**), pAKT (**c**), non-phospho-β-catenin (**d**), LC3A/B (**e**), cleaved PARP (**f**), and cleaved caspase 3 (**g**) in tumor tissues after ADT-030 vs. vehicle treatments. Welch *t*-test was used for statistical analysis. **h.** The inhibitory effect of ADT-030 on activated (GTP-bound) RAS in tumors after vehicle or ADT-030 treatments was assessed by RAS-RBD pull-down assay. **I.** Representative Ki-67 IHC results in tumors after vehicle or ADT-030 treatment. **j-l.** Representative IF images of LC3A/B (**j**), vimentin (**k**), and E-Cadherin (**l**) in tumors after vehicle or ADT-030 treatment. **m.** Bar graph representing the quantification of IHC staining for KI-67. **n-p.** Dot-plot graphs representing the IF quantifications of LC3A/B (**n**), vimentin (**o**), and E-Cadherin (**p**) in tumors after vehicle or ADT-030 treatment. Welch’s t-test was used for statistical analysis. ns, non-significant, ∗p < 0.05, ∗∗p < 0.01, and ∗∗∗∗p < 0.0001.

### ADT-030 enhances antitumor immune responses and inhibits metastasis in an orthotopic PDAC model

To confirm the anti-tumor effects of ADT-030 observed in the orthotopic 2838c3 tumors, we evaluated ADT-030 in the KPC orthotopic mouse model of PDAC to study specific immune cell subsets and functional T cell responses. To achieve this, we implanted KPC*-f-*luc cells into C57BL/6J mice, followed by treatment with ADT-030 (50, 100, or 150 mg/kg) or vehicle. Tumor growth was monitored by bioluminescence imaging on Day 0 and on Day 23 prior to euthanasia (**Fig. 5a**). The KPC model recapitulated the major findings from the 2838c3 model. Normalized bioluminescence intensities showed statistically significant differences among the vehicle and treatment groups on Day 23 (**Fig. 5b**). Both tumor size (**Fig. 5c)** and tumor weights (**Fig. 5d**) at the end of the experiment showed substantial shrinkage in a dose-dependent manner in the treated groups compared to the vehicle-treated group. We then performed IHC analysis for proliferation using the Ki-67 antibody in tumor sections from the KPC orthotopic model. Tumors treated with ADT-030 showed a strong reduction in the amount of Ki-67 positive cells compared to the vehicle group, confirming the anti-proliferative activity of ADT-030 *in vivo* (**Supplementary fig. 12a, e**). To further substantiate the above findings, we performed immunofluorescence microscopy with the tumor tissues using autophagy (LC3A/B) and the MET markers, E-cadherin and vimentin. These analyses showed a significant increase in LC3A/B, suggestive of disrupted autophagic flux (**Supplementary fig. 12b, f**), elevated E-cadherin levels (**Supplementary fig. 12c, g**), and decreased vimentin expression, reflective of MET (**Supplementary fig. 12d, h**).

**Fig. 5.**
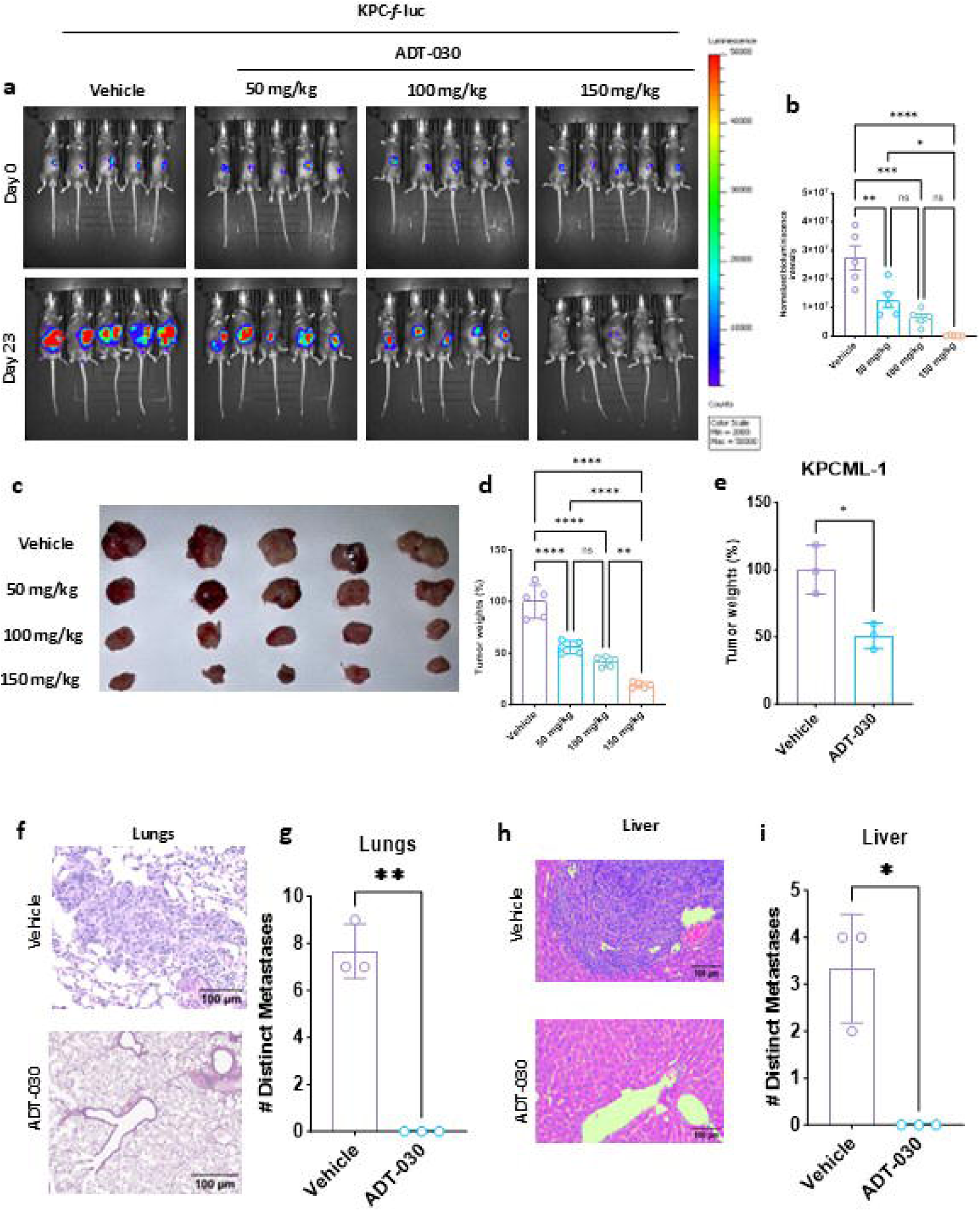
ADT-030 induces tumor growth arrest and regression in KPC and KPCML1 cells-implanted C57BL/6J mice and reduces liver and lung metastasis. **a.** KPC*-f-*luc cells were injected orthotopically into the pancreas of C57BL/6J mice. Representative bioluminescence images at the indicated time points are shown. **b.** Relative normalized whole-body bioluminescence intensities in mice under the indicated conditions (n = 5). Statistical significance was determined using one-way ANOVA. **c-d.** Tumor images after treatment with vehicle or ADT-030 at 50, 100, and 150 mg/kg (**c**) and bar graph representing tumor weights (**d**) from KPCML1 orthotopic model at termination. Statistical significance was determined using one-way ANOVA. **e.** Tumor weights from KPCML1 orthotopic model at termination. Statistical significance was determined using one-way ANOVA. **f.** Hematoxylin and eosin (HCE) staining of KPC-derived lungs after 4 weeks is shown. Representative images of HCE-stained lung metastasis sections (20x) are shown. **g.** HCE staining of KPCML1-derived liver metastasis after 4 weeks is shown. Representative images of HCE staining sections from metastasis in the lungs are displayed (20x). **h-i.** Graph representing the quantification of lung (**h**) and liver (**i**) metastatic nodules in lung sections after vehicle or ADT-030 treatment from KPC and KPCML1 orthotopic experiments. Welch *t*-test was used for statistical analysis. ns, non-significant, ∗p < 0.05, ∗∗p < 0.01, ∗∗∗p < 0.001, and ∗∗∗∗p < 0.0001

### ADT-030 promotes anti-tumor immunity in a mouse model of PDAC

We then performed multiparametric flow cytometry on the excised KPC-*f*-luc tumors to assess the immune changes underlying ADT-030 treatment. A significant elevation in overall immune infiltration was observed following ADT-030 treatment, as indicated by an increased frequency of CD45^+^ cells (**Supplementary fig. 13a**). Among the infiltrating pool of immune cells, there was an increase in αβ^+^ T cells (**Supplementary fig. 13b**), γδ^+^ T cells (**Supplementary fig. 13c**), TNK cells (**Supplementary fig. 13d**), and conventional NK cells (**Supplementary fig. 13e**), suggesting the activation of a broad-nature innate and adaptive immune response. The reduced expression of immune checkpoint molecules PD-1 (**Supplementary fig. 13f**) and CTLA-4 (**Supplementary fig. 13g**) on NK1.1^+^ cells, indicates NK cell exhaustion. In contrast, an increase in the frequencies of CD4^+^ T cells (**Supplementary fig. 13h**), as well as higher expression levels of PD-1, TIGIT, CTLA4^+^, PD-1^+^ CTLA4^+^ and FASr subsets (**Supplementary fig. 13i**), were observed in the CD4^+^ T cell compartment of ADT-030-treated tumors. The number of effector CD4^+^ T cells increased in ADT-030-treated mice, indicating involvement of helper T cell activation and functional differences (**Supplementary fig. 13j**). Similar significant increases in overall CD8+ T cell numbers were observed after ADT-030 treatment (**Supplementary fig. 13k**). In comparison, we measured decreased expression of immune checkpoint markers, including PD-1, CTLA-4, PD-1^+^CTLA-4^+^, and LAG-3, as well as lower expression of PD-1^+^CTLA-4^+^LAG-3^+^ triple-positive subsets (**Supplementary fig. 13l**) in the CD8+ T cell compartment. These data indicate a reverse of T cell exhaustion and re-engagement of cytotoxic potential. In addition, increased effector CD8^+^ T cells were observed with ADT-030 treatment, reflecting improved anti-tumor immunity (**Supplementary fig. 13m**).

Based on these results, we further explored the myeloid cell compartment in the TiME after ADT-030 treatment. We observed an enhanced influx of myeloid cells, as evidenced by increased total macrophages (F4/80^+^) (**Supplementary fig. 13n**). Another characteristic indicating myeloid infiltration was the increased expression of PD-L1 on macrophages after ADT-030 treatment, thereby enhancing antigen presentation and potential interaction with effector T cells (**Supplementary fig. 13o**). Phenotypic characterization of macrophages showed an increased M1-type characterized by MHC-II^+^CD86^+^ being more frequently expressed, while M2-like macrophages (CD206^+^) were less frequent from ADT-030 treatment compared to vehicle (**Supplementary fig. 13p-q**). Additionally, an increased M1/M2 ratio was observed after ADT-030 treatment, signifying an enhanced immune-stimulatory TiME (**Supplementary fig. 13r**). Moreover, there was also an increase in the overall frequency of dendritic cells (DCs) post ADT-030 treatment (**Supplementary fig. 13s**). Both conventional subsets of dendritic cells, cDC1, and cDC2, increased in frequency, suggesting enhanced antigen processing and presentation (**Supplementary fig. 13t-u**). This further rise in functional antigen-presenting cells, together with T and NK cell infiltration and activation, shows the wide immunomodulatory capacity of ADT-030 in remodeling of the pancreatic TiME toward an anti-tumor immune state.

### Single cell RNA-seq (scRNA-seq) of orthotopic KPC tumors treated with ADT-030

We also performed scRNA-seq on the resected tumors from this orthotopic KPC mouse experiment to further investigate the effects of ADT-030 on signaling pathways and immune microenvironment. A uniform manifold approximation and projection (UMAP) analysis was performed (**Fig. 6a**), including 10 different cellular populations identified as pericytes, gMDSCs, CAFs, endothelial cells, myocytes, T NK and B cells, dendritic cells, macrophages, acinar to ductal metaplasia (ADM), and PDAC (**Fig. 6a-b**). These groups were labelled based on the expression of recognized marker genes for mature terminal lineages (**Supplementary fig. 14a, b**). We then identified 7 PDAC sub-clusters (**Fig. 6c-d**) in which pathway analysis revealed that ADT-030-treated mice had significant downregulation in EMT, apical junction, KRAS signaling, and myogenesis signaling (**Fig. 6e**). We then focused on MAPK signaling, as this pathway is vital in driving PDAC. In ADT-030-treated mice, there was a substantial decrease in the expression of Raf1, a downstream mediator of activated RAS, suggesting the functional downregulation of the RAS-MAPK pathway in response to ADT-030 treatment (**Fig. 6f-h**). Although an increase in upstream RAS signaling was observed, the MAPK signaling flux analysis revealed a major decrease in the expression of Raf1 and Mapk3 (**Fig. 6i, j**), suggesting that downstream RAS signaling was completely inhibited. Furthermore, deep analysis revealed reduced expression of several MAPK pathway genes, including Map2k2, Mapk3, Dusp6, and Elk4 (**Fig. 6k**). We then analyzed the EMT pathway and found the concurrent decrease in the expression of mesenchymal markers for instance vimentin and fibronectin 1 (FN1) (**Fig. 6l, m**). Next, we analyzed the impact of ADT-030 treatment on WNT signaling and found that WNT pathway markers were also suppressed (**Fig. 6n**), including APC, AXIN2, Lrp5 and Lrp6 (**Fig. 6o**). Mechanistic investigation confirmed that ADT-030 treatment reduced expression of several MAPK and WNT pathway genes, supporting similar observations at the protein level (**Fig. 6p**).

**Fig. 6.**
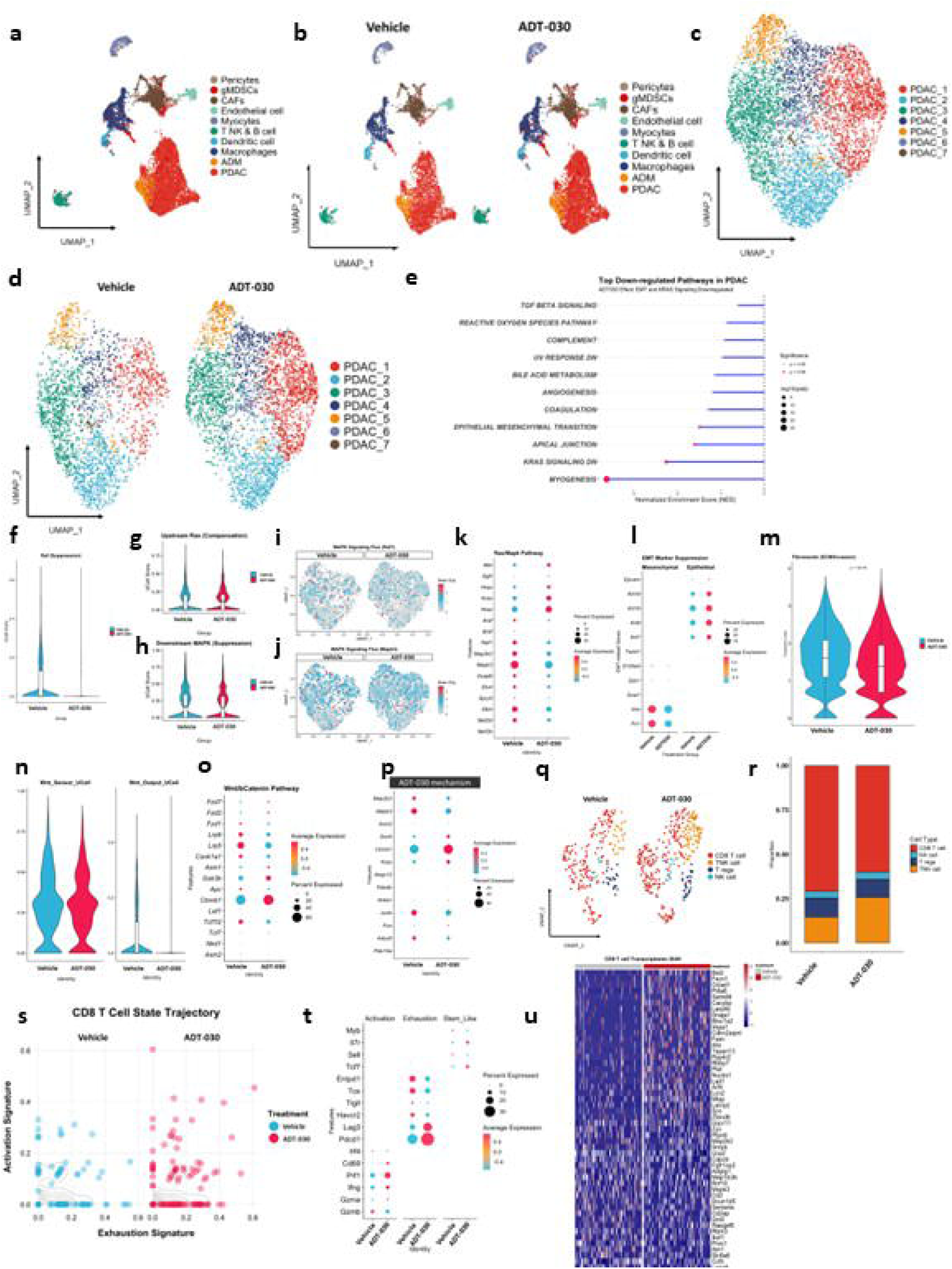
ADT-030 remodels tumor cell states and reinvigorates CD8 T cell in PDAC. **a-b.** UMAP visualization of all single cells isolated from orthotopic PDAC tumors treated with either vehicle or ADT-030, colored by major cell types as indicated. **C.** UMAP reclustering showing 7 transcriptionally distinct PDAC cell types. **d.** Split UMAPs of PDAC cells from vehicle and ADT-030 tumors demonstrating treatment-associated shifts. **e.** Bar graph showing the top significantly downregulated pathways in ADT-030-treated PDAC cells compared with vehicle controls, as determined by GSEA. **f-h.** Violin plots showing expression or module scores for RAF1 suppression (**f**), upstream RAS (**g**), downstream MAPK **(h**) related signatures in PDAC cells, including upstream WNT activation score (**f**), a downstream Wnt target gene score (**g**), and MAPK/ERK pathway score (**h**), comparing vehicle and ADT-030. **i-j.** UMAP features plots demonstrating single-cell MAPK signaling flux for Raf1 (**i**), and Map2k2 (ERK) (**j**) with corresponding dot plots summarizing average pathway activity (color scale) and fraction of PDAC cells expressing each gene set (dot size) in vehicle and ADT-030 tumors. **k-l.** Dot plots summarizing GSEA-derived pathway scores in PDAC cells, highlighting the RAS/MAPK pathway (**k**), and reduced EMT (**l**). **m-n.** Violin plots showing stemness-associated EMT-related module scores for fibronectin in PDAC cells (**m**), and WNT/β-catenin-dependent gene signatures (**n**). **o.** Dot plot of canonical WNT/β-catenin pathway genes across PDAC clusters. **p.** Dot plot of proposed mechanism-related genes illustrating transcriptional repression. **q.** UMAPs of T and NK cell compartments in vehicle and ADT-030 tumors. **r.** Stacked bar plot quantifying the proportion of CD8 T, NK, T regs and TNK cell states among total T cells in vehicle *vs*. ADT-030. **s.** Scatter plot showing CD8 T cell state trajectory scores, with each point representing a single CD8 T cell positioned according to activation and exhaustion signature scores in vehicle and ADT-030 treatments. **t.** Dot plot summarizing the expression of representative activation, exhaustion, and stem-like genes in CD8 T cells. **u.** Heat map of CD8 T cell exhaustion-associated genes across individual CD8 T cells by trajectory state comparing vehicle and ADT-030-treated tumors, depicting broad downregulation of exhaustion markers.

Next, we focused on identifying the role of ADT-030 on the immune microenvironment. To this end, we sub-clustered the UMAP into four groups: CD8 T cells, TNK cells, Tregs, and NK cells (**Fig. 6q**). A concurrent increase in the TNK cells was identified after ADT-030-treated mice, suggesting that ADT-030-treatment may enhance anti-tumor immune responses (**Fig. 6r**). Although the entire number of CD8 T cells was reduced, excessive numbers of activated CD8 T cells were present after ADT-030-treated mice compared to vehicle-treated mice (**Fig. 6s**). We also analyzed several markers of CD8 T cell activation, exhaustion, and stem-like properties. A significant increase in activation markers, including CD69, Prf1, IFNγ, and granzyme A (Gzma) was observed in ADT-030-treated mice. These findings show that ADT-030 increases CD8 T cell activation and enhances cytotoxic potential towards an effector state despite an overall reduction in total CD8 T cell numbers (**Fig. 6t**). Evaluation of the transcriptomic signature of CD8 T cells clearly showed that ADT-030 treatment increased the expression of several early-activation genes, effector differentiation factors, cytotoxicity mediators, and chemokine ligands in ADT-030-treated mice, suggesting increased CD8 T cell functionality (**Fig. 6u**).

ADT-030 treatment induced a similar but broader remodeling of the TNK compartment, extending the CD8 T-cell-specific effects to encompass both T and NK cells within the TiME (**Fig. 7a**). In line with enhanced activation and reduced dysfunction of CD8 T cells, the TNK global state trajectory showed that ADT-030 shifted TNK cells towards higher pan-activation scores and lower pan-dysfunction scores compared to vehicle treatment, indicating a coordinated reinforcement of an activated, less dysfunctional state across cytotoxic lymphocytes. This was supplemented by heightened expression and prevalence of key effector and activation genes, including Gzmb, Nkg7, and Prf1 (**Fig. 7b, c**). Additionally, ADT-030 treatment induced a marked shift in NK cell functional state toward an activated phenotype compared to vehicle treatment (**Supplementary fig. 14c**). NK cell trajectory analysis revealed that ADT-030-treated mice showed higher activation and lower dysfunction signatures, indicating coordinated enhancement of activation programs. Concordantly, dot-plot analysis revealed increased expression and prevalence of activation and maturation markers, including Zeb2, Bcl2, Klrg1, Itgam, Cd160, Havcr2, Prf1, IFNγ, and Gzmb (**Supplementary fig. 14d, e**). Together, these results demonstrate that ADT-030 enhances cytotoxic lymphocyte activation within the TiME, driving CD8 T and TNK compartments towards a sustained, less dysfunctional effector state to enhance anti-tumor immunity.

**Fig. 7.**
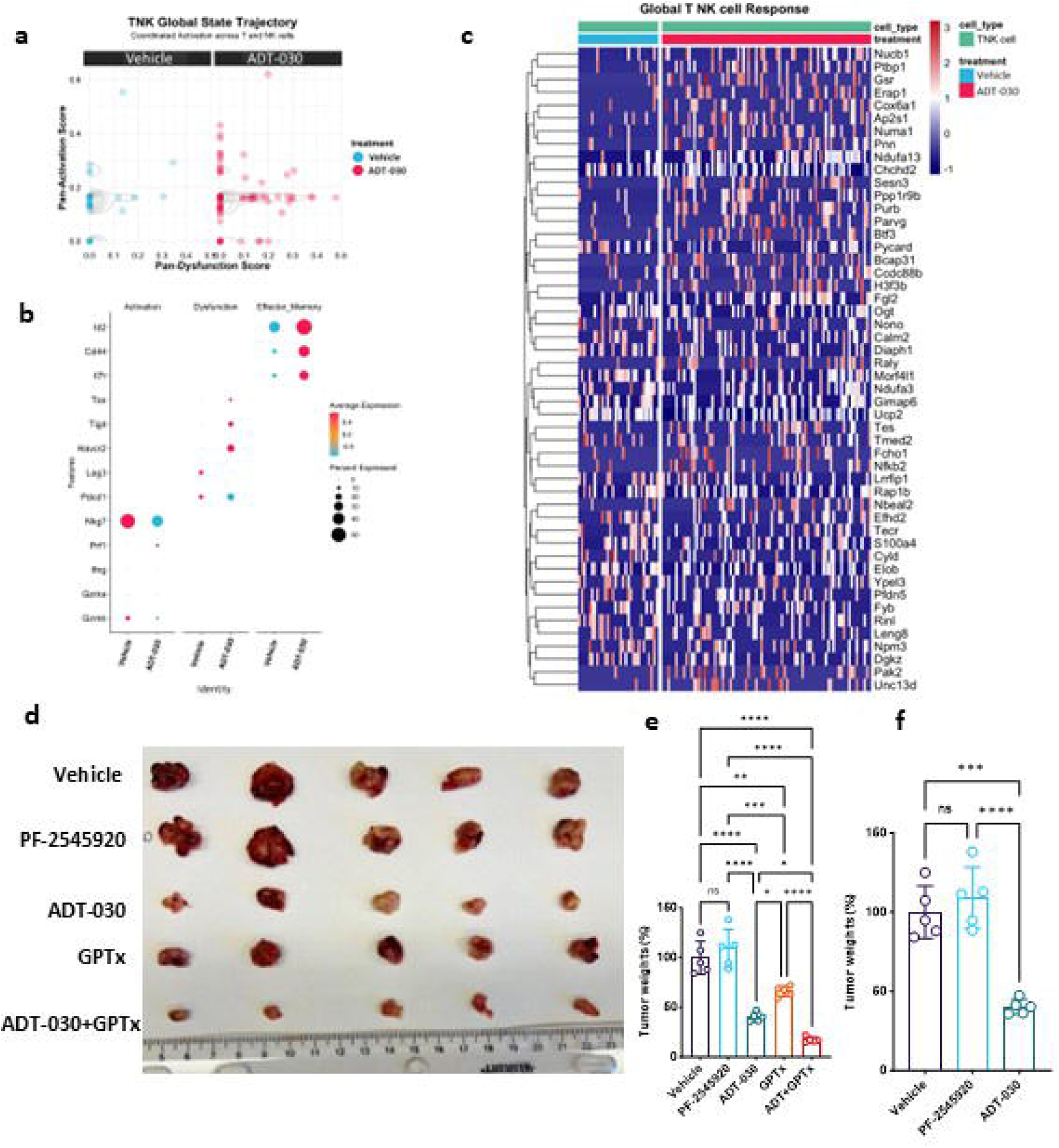
ADT-030 enhances global TNK activation and sensitizes chemotherapy to suppress PDAC tumor growth in vivo. **a.** TNK global state trajectory plot displaying pan-activation *vs*. pan-dysfunction signatures in vehicle and ADT-030 treated tumors. **b.** Dot plot showing expression of activation, dysfunction, and effector genes in TNK cells from vehicle and ADT-030 treated tumors with dot size representing the percentage of expressing cells. **c.** Heat map of differentially expressed genes in TNK cells representing a global transcriptional shift toward an activated, cytotoxic program in ADT-030-treated tumors compared to vehicle. **d.** Representative images of tumors harvested from the mice treated with vehicle, PF-2545920, ADT-030, gemcitabine paclitaxel (GPTx), and the combination of ADT-030+GPTx. **e.** Quantification of tumor weights across all treatment groups showing tumor growth inhibition with ADT-030 and further significant reduction in ADT-030+GPTx combination groups compared to monotherapies and vehicles. **f.** Comparison of tumor weights between a known PDE10 inhibitor (PF-2545920) and ADT-030 as monotherapy, demonstrating superior efficacy of ADT-030. ANOVA and Welch t-test were used for statistical analysis. ns, non-significant, ∗p < 0.05, ∗∗p < 0.01, ∗∗∗p < 0.001, and ∗∗∗∗p < 0.0001.

### ADT-030 inhibition of metastasis

To investigate the potential of ADT-030 to block metastasis, we used an established metastatic PDAC cell line, KPCML1, derived from the KPC mouse model ^36^. KPCML1 cells have a high propensity for liver and lung metastasis, as observed in patients with metastatic pancreatic cancer ^36^. Using similar tumor inoculation methods and treatment (vehicle vs. ADT-030 at 150 mg/kg daily) as described in the Materials and Methods section, we examined the impact of ADT-030 on mice orthotopically implanted with KPCML1 PDAC cells.

On day 23 after tumor implantation, ADT-030 treatment decreased the size and weight of the primary orthotopic tumor compared to vehicle treatment (**Fig. 5e**). Luciferase levels were measured using *ex vivo* imaging, which revealed that while mice in vehicle group implanted orthotopically with KPCML1 cells developed liver and lung metastasis, there was a complete absence of liver (**Supplementary fig. 15a, b**) and lung (**Supplementary fig. 15c-d**) metastasis in mice treated with ADT-030. We also analyzed lung and liver sections histologically by HCE staining, which confirmed metastasis in the vehicle group and supported the observed anti-metastatic activity of ADT-030 (**Fig. 5f-i**).

### ADT-030 enhances the antitumor efficacy of chemotherapy in orthotopic PDAC models

KPCML1 cells were implanted orthotopically and treated with vehicle, ADT-030 (150 mg/kg), standard-of-care chemotherapy (a combination of gemcitabine, 50 mg/kg and nab-paclitaxel, 10 mg/kg, GPTx), or a combination of ADT-030 with GPTx. ADT-030 produced a greater therapeutic effect than chemotherapy as indicated by tumor size and tumor weight measurements (**Fig. 7d-f**). Interestingly, the combination of ADT-030 with chemotherapy was more effective than either agent alone, suggesting that ADT-030 may enhance the efficacy of standard-of-care chemotherapy for PDAC.

### ADT-030 has increased potency and improved therapeutic window compared to conventional PDE10 inhibitors

PF-2545920 is a known PDE10 inhibitor developed for CNS disorders such as schizophrenia and Huntington’s disease. The IC_50_ values for PF-2545920 in inhibiting the proliferation of MIA PaCa-2 and 2838c cells were 25.02 and 24.1 µM, respectively (**Supplementary fig. 16a**), compared with ADT-030, which had IC_50_ values of 3.01 and 1.79 µM, respectively (**Fig. 1g**). Similarly, colony formation assays revealed that PF-2545920 had IC_50_ concentrations exceeding 25 µM (**Supplementary fig. 16b, c**), whereas ADT-030 showed significant inhibition of colony formation at 5 µM (**Fig. 1h, i**).

We then compared PF-2545920 with ADT-030 in the KPCML1 orthotopic mouse model of PDAC. Mice were treated with vehicles, ADT-030 (150 mg/kg), or PF-2545920 (10 mg/kg), each administered once daily. PF-2545920 failed to show anti-tumor activity, whereas ADT-030 significantly prevented tumor growth, as evidenced by tumor images and tumor weight measurements (**Fig. 7d-f**). Additionally, we assessed the impacts of PF-2545920 and ADT-030 treatments on mice using open-field locomotor tests. There were no noteworthy changes in behavior or mobility between ADT-030 and vehicle-treated mice, whereas PF-2545920-treated mice displayed significantly reduced mobility throughout the test, reflective of sedation, a known side effect of conventional PDE10 inhibitors (**Supplementary fig. 17a**). These data led us to conclude that ADT-030, but not PF-2545920, displays anti-tumor activity without causing sedation, which likely results from differences in brain and systemic levels between ADT-030 and known PDE10 inhibitors.

### ADT-030 suppresses tumor growth and induces prolonged responses in KRAS^G12D^ and KRAS^G12C^ PDAC PDX models

To further assess the efficiency of ADT-030 in suppressing PDAC tumor growth *in vivo*, we used two clinically annotated subcutaneously implanted PDX models of PDAC harboring KRAS^G12D^ and KRAS^G12C^ mutations. ADT-030 was administered orally at a dose of 150 mg/kg once daily for 23 days (5x/week). Tumor dimensions and body weight were measured twice/week. The results displayed a strong anti-tumor response in both KRAS^G12D^ (**Fig. 8a**) and KRAS^G12C^ PDX models (**Fig. 8d**) with no apparent systemic toxicity as evidenced by no reduction in body weight (**Fig. 8b** and **Supplementary fig. 18a, b**). The treatment was discontinued after four weeks, and mice were examined for tumor recurrence and survival. Mice treated with ADT-030 did not develop tumor regrowth over 70 days of follow-up (**Fig. 8c, f**), while vehicle-treated mice progressively died during the post-treatment period. These data demonstrate robust, durable antitumor activity of ADT-030 in clinically relevant PDAC PDX models.

**Fig. 8.**
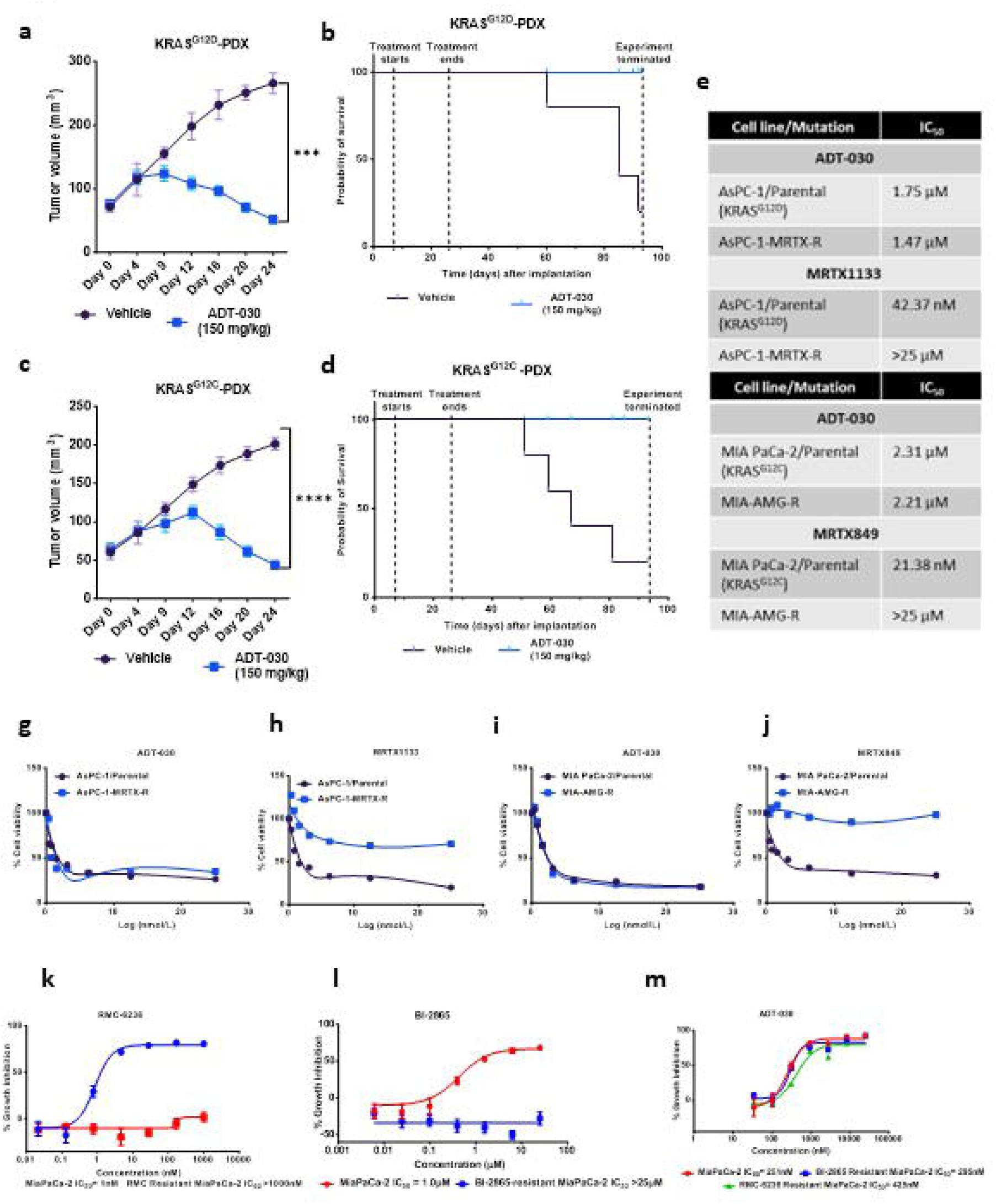
ADT-030 induces tumor regression and extends survival in KRAS^G12D^ and KRAS^G12C^ PDX models with potential to escape resistance to KRAS^G12D^ and KRAS^G12C^ inhibitors. **a.** Tumor growth in NOD.Cg-Prkdc^scid^ Il2rg^tm1Wjl^/SzJ mice implanted with KRAS^G12D^ PDX and treated with vehicle or 150 mg/kg ADT-030. **b.** Survival curves of mice after the treatment with the vehicle or ADT-030. **c.** Tumor growth in NOD.Cg-Prkdc^scid^ Il2rg^tm1Wjl^/SzJ mice implanted with KRAS^G12C^ PDX and treated with vehicle or ADT-030. **d.** Survival curves of mice after the treatment with the vehicle or ADT-030. **e.** Table showing the IC_50_ values for the cell lines used in panels. **g-j.** The indicated KRAS^G12D^-mutant PDAC cell lines were treated with various concentrations of ADT-030 (**g**) or MRTX1133 (**h**) and subjected to MTT assays. Relative percentage cell viability was plotted relative to vehicle-treated cells. **i-j.** The indicated KRAS^G12C^-mutant PDAC cell lines were treated with various concentrations of ADT-030 (**i)** or MRTX849 (**j**) and subjected to MTT assays. Relative percentage cell viability was plotted relative to DMSO-treated cells. **k-m.** Proliferation assays show that cells that are resistant to RMC-6236 (**k**) and BI-2865 (**l**) are sensitive to ADT-030 (**m**). The label on top indicates the treatment condition. Resistant cell lines were developed by exposure to sub-toxic concentrations. ∗∗∗p < 0.001, and ∗∗∗∗p < 0.0001.

### ADT-030 suppresses the growth of KRAS^G12D^ and KRAS^G12C^ - resistant PDAC cells

Given recent studies showing that Wnt activation can drive resistance to RAS inhibitors ^17^, we tested the anti-proliferative activity of ADT-030 in KRAS^G12D^- and KRAS^G12C^-resistant cell lines grown *in vitro*. AsPC-1 cells resistant to MRTX1133 (AsPC-1-MRTX-R) and parental cells were treated with ADT-030 or MRTX1133, a KRAS^G12D^ inhibitor ^37^ (**Fig. 8e-j**). Cell viability measurements using the MTT assay revealed that ADT-030 showed comparable antiproliferative activity in both parental (IC_50_ = 1.75 µM) and AsPC-1-MRTX-R cells (IC_50_ = 1.47 µM) (**Fig. 8g-h**), whereas MRTX1133 inhibited the proliferation of parental AsPC-1 cells (IC_50_ = 43.74 nM), but not of the AsPC-1-MRTX-R cells (IC_50_ > 25 µM), confirming that these cells are resistant to MRTX1133. We also determined whether this effect is sustained over longer treatment durations by determining clonogenicity. Consistent with results from proliferation assays, colony formation assays showed ADT-030 activity inhibited colony formation in both parental and AsPC-1-MRTX-R cells, while MRTX1133 inhibited colony formation only in parental cells (**Supplementary fig. 18c, d**). We also treated MIA PaCa-2^G12C^ parental and MIA PaCa-2 resistant to MRTX849 and AMG-510 (MIA-AMG-R) cells with ADT-030 and MRTX849, a KRAS^G12C^ inhibitor ^38^. ADT-030 inhibited the proliferation of both MIA PaCa-2 parental and MIA-AMG-R cells with comparable potency. In contrast, MRTX849 reduced the proliferation of MIA PaCa-2 parental cells, but not MIA-AMG-R (**Fig. 8i-j**). Similar results were found in colony formation assays, whereas ADT-030 reduced the number and size of colonies in both MIA PaCa-2 parental and MIA-AMG-R cells, while MRTX849 reduced the colony formation only in MIA PaCa-2 parental cells (**Supplementary fig. 18e, f**). Together, these results show that ADT-030 exhibits broad-spectrum RAS-inhibitory activity and has the potential to overcome acquired resistance that limits mutant-specific KRAS^G12D^ and KRAS^G12C^ inhibitors.

We also determined whether ADT-030 could overcome resistance in MIA PaCa-2 PDAC cell lines resistant to the pan-RAS inhibitor RMC-6236 and the pan-KRAS inhibitor BI-2865 (**Fig. 8k-m**). Results indicated that both resistant cell lines were completely not sensitive to RMC-6236 and BI-2865 but remained fully sensitive to ADT-030, with IC_50_ values comparable to those of the parental cells (**Fig. 8k-m**). These findings suggest that ADT-030 remains active even after resistance to pan-RAS inhibitors.

### Effects of long-term ADT-030 exposure on metabolic reprogramming

To investigate the mechanism underlying adaptive resistance to PDE10 inhibition, we performed long duration incubation of PDAC cell model with ADT-030. We found that PDE10 expression was markedly downregulated in these cells (**Supplementary fig. 19a**), indicating suppression of PDE10 as a potential mechanism of adaptive resistance. To characterize this adaptive phenotype, we employed a Human Central Metabolism Array encompassing 14 pathways. Our findings indicate a hybrid metabolic program in ADT-030-resistant cells, marked by synchronized upregulation of genes associated with glycolysis, fatty acid oxidation, and nucleotide biosynthesis (**Supplementary fig. 19b-e** and **Supplementary fig. 20a-h**). We also observed an elevated expression of essential glycolysis regulators (*SLC2A5*, *SLC2A14* (*GLUT14*), and *PFKFB3*) and nucleotide metabolism enzymes (*RRM1*). ADT-030-resistant cells showed more general compensatory rewiring. This included the activation of *ACADL*, *CPT2*, *EHHADH*, and reduced expression of GLS isoforms, as well as the adenylate kinase (AK) family (*AK3* and *AK8*). These findings collectively indicate that continuous exposure to ADT-030 induces a stable, metabolically reprogrammed state that facilitates survival by concurrently activating glucose, lipid, and amino acid metabolism.

## Discussion

Aberrant activation of MAPK/AKT signaling from KRAS mutations, along with the stimulation of WNT/β-catenin-mediated transcription, plays a major role in driving cancer cell proliferation, survival, and metastasis in PDAC and other cancers, including colorectal, liver, lung, and breast cancer ^39–42^. KRAS is mutated in over 90% of PDAC, while mutations or increased expression of β-catenin or other pathway components (e.g., Wnt ligands, FZD receptors, LRP co-receptor, APC, Axin) are also observed in a high percentage of patients with PDAC and other cancers ^43,44^. Compensatory or cooperative interactions between these signaling pathways likely also contribute to the aggressiveness of the disease, as well as therapy resistance ^45,46^.

Phosphodiesterase isozymes have been previously studied in the perspective of cancer, but no particular isozyme has been targeted by an inhibitor, and no PDE inhibitor has achieved FDA authorization for the treatment of cancer ^23^. Lately, numerous publications have reported that isozyme-specific PDE10 inhibitors and genetic silencing of PDE10 can principally hinder the proliferation and persuade apoptosis of colon, lung, and ovarian cancer cell lines expressing PDE10 by activating cGMP-dependent PKG to block RAS and β-catenin signaling, but not cells from normal tissues lacking PDE10 expression ^27–30^. cAMP-dependent-PKA has been reported to inhibit signaling downstream of RAS by disrupting interaction with Raf1 ^47^. Our data revealed increased expression of PDE10, a dual cGMP and cAMP-degrading PDE isozyme, in PDAC cells compared with adjacent normal pancreatic tissue, providing an initial rationale for targeting this pathway in PDAC treatment. Additionally, we found that PDE10 mRNA expression is positively correlated with KRAS mRNA expression, suggesting that both oncoproteins and their respective signaling pathways cooperate to drive PDAC progression. These conclusions are supported by the aforementioned studies, which report PDE10 overexpression in other cancers and an association with reduced patient survival ^25,27–30,48^.

ADT-030 is a non-COX inhibitory sulindac derivative and a second-generation analog of ADT-061 (aka MCI-030), previously stated to selectively impede the PDE10 isozyme at concentrations that inhibit the proliferation and induce apoptosis of colorectal and ovarian cancer cell lines ^27,30^. Previous studies reported that PDE10 knockdown had RAS-selective antiproliferative and pro-apoptotic activity and reduced the sensitivity of cancer cells to the growth-inhibitory activity of ADT-061, as well as known PDE10 inhibitors, confirming the selectivity of this class of agents for PDE10 and that PDE10 is important for cancer cell growth and survival ^27–30^. Similarly, in our study, we found that PDE10 knockdown cells are more vulnerable to ADT-030’s growth-inhibitory activity than parental cells, further supporting the notion that PDE10 is the primary molecular target of ADT-030. Our findings in PDAC cells are supported by other investigators also reporting the role of PDE10 in inflammation or cancer ^48–50^. Molecular docking simulations and cellular engagement thermal stability assays presented in this study provide structural insight into the interaction between ADT-030 and PDE10 and confirmation of target engagement, respectively. Together, these results deliver additional evidence for an oncogenic role of PDE10 and support a mechanism of action for ADT-030 involving PDE10 inhibition, elevated cyclic nucleotide levels, and protein kinase activation.

Of translational significance, we found that ADT-030 prevents the proliferation of PDAC cells that develop resistance to mutant-specific KRAS and pan-RAS inhibitors, suggesting its potential to overcome a major clinical obstacle for RAS inhibitors that are approved or in development. The potential to escape resistance may been attributed to the capacity of ADT-030 to induce cell cycle arrest and apoptosis, resulting from the dual blockage of β-catenin and RAS signaling. The observed inhibition of MAPK and AKT/PI3K pathways by ADT-030 is particularly significant for PDAC treatment, as both pathways are known for their extensive crosstalk and compensatory activation to drive cancer cell proliferation and survival ^51,52^. Compensation from β-catenin may also contribute to resistance to monospecific inhibitors of KRAS or β-catenin, whereby dual blockage of RAS or β-catenin pathways through PDE10 inhibition may prevent the KRAS inhibitor resistance, FDA-approved or in development ^53^. To corroborate these findings using gene expression profiling, we evaluated tumors excised from mice treated with ADT-030 or vehicle using single-cell transcriptomics. The results confirmed suppressive effects of ADT-030 on key oncogenic signaling pathways, including RAS-MAPK, EMT, and WNT, as evidenced by reduced expression of Raf1, Mapk3, Map2K2, vimentin, FN1, APC, and Axin2. We also conducted assays on RAS activation in RAS wild-type and KRAS mutant PDAC cell lines and found that ADT-030 selectively inhibited RAS activation in KRAS mutant PDAC cells. This interesting observation needs further study to understand the differential effects of PDE10 inhibition and the impact of cyclic nucleotide signaling in KRAS mutant PDAC cells compared with RAS wild-type PDAC cells.

ADT-030 is orally bioavailable with attractive drug-like properties and appears to be well tolerated at dosages that exhibit robust and durable antitumor activity. We found that ADT-030 inhibits both primary tumor growth and metastasis without discernible toxicity in several mouse models of PDAC, including PDX and orthotopic models. Of clinical significance, ADT-030 also potentiated the efficacy of standard-of-care chemotherapy regimens for PDAC. These findings encourage the rationale for developing ADT-030 as a front-line treatment for PDAC, either as monotherapy or in combination with standard-of-care chemotherapy.

PDE10 inhibitors have been previously developed for the treatment of CNS disorders such as schizophrenia and Huntington’s disease. We therefore compared ADT-030 with the known PDE10 inhibitor PF-2545920 and found that ADT-030 exhibited appreciably greater potency than PF-2545920 at inhibiting PDAC cell proliferation *in vitro*. This observation suggests that although PDE10 is a cancer target, PF-2545920 has low potency to inhibit cancer cell proliferation, which may be attributed to compensation by other co-expressed PDE isozymes (e.g., PDE5). Compared with ADT-030 in vivo, PF-2545920 failed to demonstrate anti-tumor activity. In addition, ADT-030 displayed improved tolerance without the side effects (sedation) observed with PF-2545920 ^54^.

PDAC is also associated with an immunosuppressive TiME, a major factor responsible for resistance to immunotherapy ^55,56^. The immune infiltration in PDAC is portrayed by the abundance of immune suppressive cells and the lack of anti-tumor immune cells ^57^. Activation of the immune system by ADT-030 treatment is another intriguing and clinically relevant finding reported here. ADT-030 treatment increased CD4^+^ and CD8^+^ T cells, as well as NK cell infiltration, leading to a shift towards M1-like macrophage polarization. The inhibition of expression of CTLA-4, PD-1, and LAG-3 on CD8^+^ T cells by ADT-030 treatment suggests that ADT-030 alleviates T cell exhaustion and reestablishes cytotoxic T cell function within the TiME ^58^. Aside from maintaining antitumor T cells, ADT-030 also enhanced myeloid cell infiltration by increasing the number of total macrophages (F4/80^+^) within the TiME. These macrophages display enhanced antigen-presenting capacity, as evidenced by increased PD-L1 expression, which is crucial for effector T cell interaction. In addition, phenotypic analysis revealed a shift in macrophage polarization toward a pro-inflammatory, M1-like phenotype (MHCII^+^ CD86^+^), accompanied by a reduction in M2-like macrophages (CD206^+^) ^59^. These results were further supported by the identification of a high M1/M2 ratio. In addition to macrophages, ADT-030 treatment increased conventional dendritic cells (cDC1 and cDC2), thereby contributing to the activation of T and NK cells ^60^. Our single-cell RNA sequencing data demonstrated that the cytotoxic lymphocyte compartment is reprogrammed, with CD8 T cells and TNK cells exhibiting enhanced activation, effector function, and reinvigoration. Activation of cytotoxic genes (IFNγ, Gzma, and Prf1) and activation markers (CD69 and Cxcr3), along with modulation of exhaustion pathways (Lag3 and CTLA4), indicates that ADT-030 not only suppresses tumor cell proliferation but also potentiates anti-tumor immunity. These convergent tumor and immune-cell effects provide a strong mechanistic rationale for evaluating ADT-030 in combination with immune checkpoint blockade or other immunomodulatory strategies to convert immunologically cold PDAC into a more treatment-responsive state. Interestingly, this will be a future direction for preclinical studies combining ADT-030 with immune checkpoint blockade, as well as for possible clinical trials, given the limitations of immunotherapy for PDAC.

While mutant-selective KRAS^G12C^ and KRAS^G12D^ inhibitors have demonstrated promising efficacy for KRAS mutant cancers ^61,62^. Acquired resistance remains a major clinical limitation. Several resistance mechanisms have been reported, comprising secondary RAS mutations, activation of co-expressed RAS wild-type isozymes, and compensatory receptor tyrosine kinase mutations, all of which frequently emerge in recurrent tumors and contribute to treatment failure, disease relapse, and death of the patient ^63^. In the current study, we investigated whether ADT-030 may be less susceptible to the same mechanisms of resistance that limit the efficacy of KRAS^G12C^ and KRAS^G12D^ inhibitors using PDAC cell lines developed to be resistant to such drugs. Strikingly, ADT-030 demonstrated potent anti-proliferative activity in both MRTX1133- and MRTX849-resistant PDAC cell lines, highlighting its broad-spectrum pan-RAS inhibitory activity and its ability to bypass diverse mechanisms of acquired resistance to allele-specific KRAS inhibitors.

The therapeutic potential of ADT-030 is supported by antitumor results observed in clinically and genetically relevant PDAC PDX models harboring KRAS^G12D^ and KRAS^G12C^ mutations as well as several orthotopic mouse models. In these experiments, the efficacy and tolerability of ADT-030 were assessed following oral administration at 150 mg/kg for 4 weeks. This dosage caused no discernible toxicity and is a human equivalent dosage of 12 mg/kg, or 840 mg once daily for a 70 kg human, a moderate dose for many drugs. Treated mice showed tumor regression and no tumor regrowth for over 70 days after stopping treatment, highlighting the durability in maintaining long-term tumor control. These findings provide compelling evidence of robust, clinically relevant antitumor activity of ADT-030 by inhibiting PDE10, supporting IND-enabling studies. This activity and ADT-030’s capacity to simultaneously block RAS and β-catenin signaling also support further mechanistic studies of the oncogenic role of PDE10 (**Supplementary fig. 21**). Future studies focusing on the activity of ADT-030 in genetically engineered mouse models, both as monotherapy and in combination with immune checkpoint inhibitors, will further help translate this agent to the clinic. In conclusion, the ability of ADT-030 to inhibit PDE10 and block multiple aspects of malignant progression, including cancer cell proliferation, survival, and metastasis, as well as creating a more favorable TiME, while having the potential to escape resistance that limits the efficacy of other RAS inhibitors, makes ADT-030 a highly desirable drug development candidate for clinical trials in patients with metastatic PDAC and other RAS-driven malignancies.

## Materials and methods

### ADT-030 synthesis

The synthesis of ADT-030 was performed as reported in our previous publication ^64^.

### Human scRNA-seq data mining

The Single Cell RNA seq Pancreatic Cancer Atlas R Data Serialization (RDS) file (https://zenodo.org/records/14199536) was downloaded ^65^. This dataset contains normalized and scaled scRNA-seq (10x Genomics) data from 12 studies, including 229 patients across groups. The ductal cells were identified from the main dataset, and the expression levels of PDE10 were queried across the tissue samples (donor, adjacent normal, primary tumor, and metastatic lesion) as described in the previously published paper^65^.

### Cellular thermal stability assay

HEK293 cells expressing PDE10 fused to the MICRO-TAG reporter were subjected to a temperature-series denaturation assay to determine the aggregation midpoint under cellular conditions, as previously described ^66^. Cells were heated for 10 min across a defined temperature range, followed by non-denaturing lysis and fluorescence complementation quantification. Fitting the resulting thermal curve yielded a Tagg₅₀ of 44°C for PDE10. This defined Tagg₅₀ served as the fixed challenge temperature for subsequent experiments to determine whether ADT-030 binds PDE10 in intact cells.

### Cell culture

PANC-1 (ATCC# CRL-1469; RRID:CVCL_0Q68), AsPC-1 (ATCC# CRL-1682; RRID:CVCL_0152), Panc 02.03 (ATCC# CRL-2553; RRID:CVCL_1633), Panc 10.05 (ATCC# CRL-2547; RRID:CVCL_1639), MIA PaCa-2 (ATCC# CRL-1420; RRID:CVCL_0428), BxPC-3 (ATCC# CRL-1687; RRID:CVCL_0186), and KLE (ATCC# CRL-1622; RRID:CVCL_1329) cell lines were obtained from American Type Culture Collection (ATCC, Manassas, VA, USA). Mouse PDAC cell line 2838c3 (Kerafast# EUP013-FP; RRID:CVCL_YM18) was purchased from Kerafast (Boston, MA, USA). MKN1 (Accegen# ABC-TC0685; RRID:CVCL_1415) cell line was purchased from Accegen Inc (Fairfield, NJ, USA). Dr Gregory Lesinski of Emory University, USA, gifted the KPC cell line. Dr. Denis C Guttridge, Medical University of South Carolina, USA, gifted the KPCML1 cell line. All cells were grown in appropriate medium as recommended by the ATCC and Kerafast. AsPC-1 and MIA-PaCa-2 cells resistant to MRTX1133 and MRTX849 were developed and maintained as described previously ^67,68^. PANC-1 cells resistant to ADT-030 were generated by continuous exposure to ADT-030 for a period of 31 days with increasing doses starting from the IC_50_ concentration. The maximum resistance dose achieved was 10 µM. MIA PaCa-2 cells were persistently dosed to RMC-6236 or BI-2865 starting at their respective IC_50_ concentrations. Drug concentrations were stepwise increased as cells recovered and confluency increased. Resistant populations were established and maintained at 100nM (RMC-6236) or 2.5µM (BI-2865).

### Phosphodiesterase assay

The enzymatic activity of recombinant PDE10 was measured using the Immobilized Metal Affinity Particle (IMAPTM) fluorescence polarization (FP) progressive binding system (Molecular Devices; San Jose, CA; USA) as earlier described to determine the inhibitory effect of ADT-030 ^69^. FP was assessed using a Synergy H4 Hybrid plate reader (BioTek; Santa Clara, CA; USA). Recombinant PDE10 was procured from BPS Biosciences (San Diego, CA; USA).

### PDE10 KD stable cell line preparations

Human PDE10-specific and non-silencing (NS) control short hairpin RNA (shRNAs) were acquired from Sigma Aldrich. Human PDAC cells (PANC-1 and AsPC-1) were transduced with these lentiviral particles and selected with puromycin at 0.75 µg/ ml. Total cell lysate was isolated from the selected cells, and immunoblotting was performed to validate the knockdown. β-actin was used as a loading control.

### Proliferation assay

PDAC cells with KRAS mutations (KRAS^G12D^ and KRAS^G12C^) and wild-type cells (BxPC-3) were used to test the proliferation assays as reported in our previous publication ^70,71^.

### Clonogenic assay

Mouse-derived PDAC cell line 2838c3 (KRASG12D) and human-derived cell line MIA PaCa-2 (KRAS^G12C^) were seeded in 6-well culture plates (2 × 10^3^ cells/well), and the experiment was performed as reported in our previous studies ^70,71^.

### Motility assay

To assess the impact on cell motility, 2838c3 and MIA PaCa-2 cells were cultured in 6-well plates until they reached near confluence. The cells received treatment with either DMSO or ADT-030 at concentrations of 0.5, 1, 2, and 5 µM. A mark was made with a sterile 10-µL pipette tip, and cell migration was observed each day under light microscopy. Cell movement was quantified using ImageJ.

### Apoptosis assay

The binding of annexin V to cells was determined with the PE-Annexin V Apoptosis Detection Kit I (BD Biosciences# 559763), according to our previously established protocol ^71^.

### Cell cycle assay

2838c3 and MIA PaCa-2 cells were treated with either DMSO or ADT-030 (2 and 5 µM) for 24 hrs and the experiment was performed as reported in our previous studies ^71^.

### Total RNA isolation, cDNA preparation, and RT-PCR

Total RNA was isolated from ADT-030-resistant and parental cells using TRIzol reagent (Invitrogen, Carlsbad, CA, USA) and refined with an RNeasy Mini Kit (Qiagen). cDNA was synthesized using the First Strand cDNA Synthesis Kit (Arraystar #AS-FS-001) as per the manufacturer’s instructions. Next, qPCR was performed using the human central metabolism PCR array (Arraystar #AS-NM-004), and the data were analysed per the manufacturer’s guidelines.

### Immunoblotting

Immunoblotting was done as reported in our previous studies ^70,71^. Primary antibodies were pERK, ERK, pAKT, AKT, pmTOR, mTOR, pP70s6 kinase, p70s6 kinase, pCREB, CREB, Bcl-2, VEGFA, PDE3B, PDE4C, PDE4D, LC3A/B, cleaved PARP, cleaved caspase 3, non phospho β-catenin, β-catenin, pVASP, VASP, and anti-β-actin. Incubation with HRP-linked secondary antibodies (CST, #7074/7076) at a dilution of 1:3000 in 2% BSA was carried out for 1 hr at room temperature. Additionally, the PANC-1 and BxPC-3 cells were treated with either DMSO or 5 µM ADT-030 for 24 hrs, to isolate total lysates. The lysates were then probed to detect metabolic changes using the Human Metabolic Pathways Array C2 (RayBiotec # AAH-META-2) fitting to the manufacturer’s information.

### Measurement of intracellular cAMP and cGMP levels

PDAC cell lines, 2838c3 and MIA PaCa-2, were treated with ADT-030 at varying doses, harvested, and the intracellular cAMP and cGMP levels were measured by ELISA. Enzyme immunoassay kits were used to detect cAMP (Cat# 581001, Cayman) and cGMP (Cat# 581021, Cayman) by following the manufacturer’s protocol. The results were defined as picomoles/µg of total protein.

### ELISpot

Tumors obtained from the orthotopic models containing 2838c3 cells were mechanically homogenized using a syringe plunger, passed through a 40-μm cell strainer, and erythrocytes were lysed with a RBC lysis. Freshly isolated single cells (2 × 10⁵ cells/well) were then co-cultured with each neoantigen peptide (10 ng/µL) in RPMI-1640 medium enriched with 10% FBS and incubated in IFN-γ ELISpot plates (Mabtech, USA; 3321-4APT-2) for 18–24 hours at 37 °C. Negative controls were resuspended in RPMI-1640 medium with 10% FBS. The T cells secreting IFN-γ were subsequently detected by IFN-γ ELISpot assays corresponding to the manufacturer’s guidelines. The subsequent spots were quantified utilizing ImageJ.

### Immunohistochemistry (IHC) and immunofluorescence (IF)

Paraffin-embedded tumor tissue slides from 2838c3 and KPC orthotopic studies were used to detect the expression patterns of extracellular matrix (ECM) remodeling, apoptosis, and autophagy markers by IHC and IF by following the established protocol from our previous studies ^70,71^.

### Active RAS detection assay

Active RAS (RAS-GTP) levels were assessed by the Active RAS Activation Assay Kit (Cell Signaling Technology, Cat# 8821) by following the established protocol from our previous publication ^70^.

### Pharmacokinetics, tissue distribution, and histopathological examination of ADT-030 in mice

Pathogen-free 8-week-old female C57BL/6J mice (Envigo#044; RRID:IMSR_ENV:HSD-044) were housed in the Biologic Research Laboratory at the University of South Alabama (USA), Frederick P. Whiddon College of Medicine with *ad libitum* access to food and water throughout the experiments. Following acclimatization, mice were treated with ADT-030 at a dose of 100 mg/kg once daily for 14 days by oral gavage. After the last administration, blood was drawn and collected into K2EDTA Microtainer tubes (BD Biosciences; Franklin Lakes, NJ, USA) at 0.5, 1, 2, 4, 8, and 24 hrs (n=4 per time point) to obtain plasma. Major organs (lungs, kidneys, spleen, heart, liver, brain, colon, and ovaries) were collected at 8 hrs (n=4). LC-MS/MS determined ADT-030 levels in plasma and organs. The study followed established guidelines and adhered to the approved protocol of the USA Institutional Animal Care and Use Committee (IACUC).

In another study, following adaptation at the UAB animal facility, 5-6-week-old male C57BL/6J mice (The Jackson Laboratory #000664; RRID:IMSR_JAX:000664) were randomly assigned to two groups (n=5 per group) and received ADT-030 (150 mg/kg) by oral gavage once daily for 2 weeks or vehicle. At the end of the treatment, blood was drawn for serum biochemical analysis followed by mice euthanize for organ collection and bone marrow smears preparation for histopathological and cytology analysis, respectively. Following standard procedures, a board-certified veterinary anatomic pathologist (JBF) performed a blinded assessment of visceral organs (heart, lung, kidney, liver, duodenum, pancreas, colon, spleen, thymus, testes) and brain.

### Open field locomotor activity

Mice had *ad libitum* access to food and water throughout the experiments. Behavioral experiments were performed during the light cycle (between 8 a.m. and 6 p.m.). Before evaluation, mice were housed in the testing room for at least 30 min. Mice were placed in an open-field arena (44×44×30 cm) in a dimly lit room (7 lx) and allowed to freely explore for 10 min as previously described ^72^. Locomotor activity was identified as the cumulative distance traveled during the entire 10 min. All behavioral data was acquired using Noldus EthovisionXT (Version 16) to track and record mice. Experimenters were blinded to treatment for all comparisons between groups.

### Orthotopic grafting of PDAC cells in mice

*In vivo* studies followed established guidelines and adhered to the approved UAB IACUC protocol. All the studies with indicated cells lines has been perfomed as per the established protocol from our previous publications ^70,71^. Mice in PKC and 2838c3 studies were given oral dosages as follows: the first group received a vehicle, and the other three groups received ADT-030 at varying oral dosages (50, 100, and 150 mg/kg). In the KPCM1 study, mice received vehicles or ADT-030 at 150 mg/kg. In the combination study with KPCML1 cells implanted, mice received vehicle, ADT-030 (150 mg/kg), or PF-2545920 (10 mg/kg, IP) daily for 4 weeks, gemcitabine (50 mg/kg, IP) plus nab-paclitaxel (30 mg/kg, IP), twice weekly (GPTx), and ADT-030 plus GPTx. Tumor tracking and response to therapy were monitored using D-luciferin injections and were performed bi-weekly through the studies. The whole luminescence from tumor-bearing zones was measured using the Living Image *in vivo* imaging software. The animals’ body weights were calculated twice a week. At the end of the study, each animal underwent whole-body imaging, after which they were euthanized. During this process, tumors were harvested, weighed, and utilized for further experiments.

### Single-cell tumor processing

FFPE blocks of the tumor tissues from KPC orthotopic experiments treated with vehicle or ADT-030 (150 mg/kg) were used for sc-RNA sequencing, as per the standard protocol used at Admera Health (South Plainfield, NJ, USA). After sequencing, the data were analyzed by demultiplexing and aligned to the mouse reference genome (GRCm39) for gene expression quantification and processed with Cell Ranger 9.0.1. The count matrices were then analyzed in the Seurat (v.5.3.0) R package (v.4.5.1). Cells with more than 200 genes and less than 5% mitochondrial content were kept for downstream analysis. After SC Transform analysis, PCA and UMAP were used for dimensionality adjustment and clustering. Differentially expressed genes (DEGs) were identified using FindAllMarkers, and clusters were labeled using known markers.

### Tumor growth inhibition studies using PDAC PDX tumors

Mouse experiments were conducted using PDAC PDX models with KRAS^G12D^ and KRAS^G12C^ mutations as described previously ^70^. The first group received vehicle, and the second group received ADT-030 (150 mg/kg, 5x/day). A survival study was conducted for 70 days, beginning 28 days after treatment.

### Flow cytometry

Tumors derived from 2838c3*-f-*luc and KPC*-f-*luc PDAC cells implanted into the pancreas of C57BL/6J mice were used to identify the changes in immune phenotype as per the established protocol from our previous publications ^70,71^. The antibodies and dilutions were mentioned in (**Supplementary table 1**). Additionally, single cells were used to perform ELISpot to detect IFN-γ expression (MABTECH #3321-4APT) following manufacturer’s instruction.

### Statistical analysis

Statistical analyses and data visualization were done using GraphPad Prism (RRID:SCR_005375). The data are represented as means accompanied by either standard deviation (SD) or standard error of the mean (SEM). A repeated-measures analysis of variance (ANOVA) or an ANOVA with a Bonferroni correction was conducted to assess statistical significance between groups. A statistical significance threshold was set at p < 0.05.

#### Data and materials availability

All data associated with this study are presented in the main publication or supplementary figures. All materials used to conduct the experiments will be shared upon request, subject to a suitable agreement. All code used to process the scRNA-seq data and to draw the figures is available on GitHub at the following link: https://github.com/sujithsrvsh/ADT030_scRNAseq, and the sequence is available under the following accession number: GSE327617.

## Acknowledgments

This work was supported by the University of Alabama at Birmingham (UAB), Birmingham, AL, USA. B.E was supported by the UAB O’Neal Comprehensive Cancer Center, National Institutes of Health (NIH) core support grant 5P30CAO13148-47 and 1R01CA294647. This work was also funded by the NIH grants R01CA254197 (Piazza) and R01CA238514 (Zhou and Piazza).

## Conflict of interest

A.B.K, X.C., and G.A.P are assignees on patents covering compositions of matter for ADT-030 and analogs. All other authors declare no competing interests.

## Contributions

Concept and design: BE and GP; acquisition, analysis, or interpretation of data: DSRB, VRA, GSG, LC; drafting of the manuscript: DSRB; critical revision of the manuscript for important intellectual content: all authors; statistical analysis: DSRB; administrative, technical, or material support: BE; validation: DSRB; supervision: BE. sc-RNA data analysis: SS; Modeling: TH; Histopath: JBF; Thermal assay: EN and IB; In vitro testing: ABK; All authors have read and approved the article.

## Notes

### Competing Interest Statement

A.B.K, X.C., and G.A.P are affiliated with ADT Pharmaceuticals, LLC. All the other authors declare no competing interest

### Summary of Updates

We have revised the manuscript per the reviewers' comments, and this is the most recent version before acceptance.

